# Di- and tri-methylation of histone H3K36 play distinct roles in DNA double-strand break repair

**DOI:** 10.1101/2023.10.18.562911

**Authors:** Runfa Chen, Meng-Jie Zhao, Yu-Min Li, Ao-Hui Liu, Ru-Xin Wang, Yu-Chao Mei, Xuefeng Chen, Hai-Ning Du

## Abstract

Histone H3 Lys36 (H3K36) methylation and its associated modifiers are crucial for DNA double-strand break (DSB) repair, but the mechanism governing whether and how different H3K36 methylation forms impact repair pathways is unclear. Here, we unveil the distinct roles of H3K36 dimethylation (H3K36me2) and H3K36 trimethylation (H3K36me3) in DSB repair via non-homologous end joining (NHEJ) or homologous recombination (HR). Yeast cells lacking H3K36me2 or H3K36me3 exhibit reduced NHEJ or HR efficiency. yKu70 and Rfa1 bind H3K36me2- or H3K36me3-modified peptides and chromatin, respectively. Disrupting these interactions impairs yKu70 and Rfa1 recruitment to damaged H3K36me2- or H3K36me3-rich loci, increasing DNA damage sensitivity and decreasing repair efficiency. Conversely, H3K36me2-enriched intergenic regions and H3K36me3-enriched gene bodies independently recruit yKu70 or Rfa1 under DSB stress. Importantly, human KU70 and RPA1, the homologs of yKu70 and Rfa1, exclusively associate with H3K36me2 and H3K36me3 in a conserved manner. These findings provide valuable insights into how H3K36me2 and H3K36me3 regulate distinct DSB repair pathways, highlighting H3K36 methylation as a critical element in the choice of DSB repair pathway.

## Introduction

DNA double-strand breaks (DSBs) are highly cytotoxic DNA lesions that cause genomic instability and detrimental chromosomal rearrangements (Kockler *et al*., 2021, Symington & Gautier, 2011), potentially leading to severe human diseases like cancer and immunodeficiency responses (Chiruvella *et al*., 2013, Kowalczykowski, 2015). The repair of DSBs mainly occurs through two pathways: non-homologous end joining (NHEJ) and homologous recombination (HR), both operating within chromatin to safeguard the genome (Kowalczykowski, 2015, Li *et al*., 2019). NHEJ restores chromosome integrity by directly joining broken DNA ends, relying on the Ku heterodimer complex (Ku70/Ku80) to prevent DNA degradation and recruit other NHEJ factors (Daley *et al*., 2005, Fell & Schild-Poulter, 2015). Conversely, HR utilizes the replication protein (RPA) complex to coat single-stranded DNA, facilitating HR repair through Rad51 nucleofilament formation (Jasin & Rothstein, 2013, Marechal & Zou, 2015, Symington, 2014, Symington & Gautier, 2011). Failure or misuse of each pathway may result in deleterious consequences on the genome. The selection between these pathways involves several factors, including cell cycle stage, DNA end structure, and epigenetic context (Findlay *et al*., 2018). Although the landscape of chromatin marks near DSBs has been studied, the specific roles of individual histone modifications in regulating DSB repair pathways remain largely unknown.

A plethora of post-translational modifications on histones participate in DNA damage response (DDR) and DNA repair (Bird *et al*., 2002, Iacovoni *et al*., 2010, Peng *et al*., 2021b, Smeenk & van Attikum, 2013). Among these modifications, histone H3 lysine 36 methylation (H3K36me) stands out as a critical mark that is deposited by various methyltransferases in different species (Wagner & Carpenter, 2012). In *Saccharomyces cerevisiae* (hereafter referred to as budding yeast), the methyltransferase Set2 is responsible for all three mono-, di-, and tri-methylation of H3K36, abbreviated as H3K36me1, H3K36me2, and H3K36me3, respectively. Conversely, in humans, NSD1, NSD2/MMSET, NSD3, and ASH1L are involved in catalyzing H3K36me1 and H3K36me2, while SETD2 specifically adds H3K36me3 (Husmann & Gozani, 2019). The roles of H3K36 methylation are conserved from yeast to humans and are associated with various fundamental biological processes, including transcription regulation, transcription fidelity, RNA processing, DNA replication, recombination, and repair (Dronamraju *et al*., 2018, McDaniel & Strahl, 2017, Pai *et al*., 2017, Pfister *et al*., 2014, Wagner & Carpenter, 2012). For DSB repair, studies have suggested that H3K36me2 promotes NHEJ in mammals (de Krijger *et al*., 2020, Fnu *et al*., 2011), while SETD2-catalyzed H3K36me3 facilitates HR repair by aiding DNA resection and promoting RAD51 filament formation (Pfister *et al*., 2014). In yeast, cells lacking Set2 show impaired activation of the DNA-damage checkpoint and weakened DSB repair (Jha & Strahl, 2014). However, there have been conflicting findings in fission yeast, where Set2-dependent H3K36me3 seems to promote NHEJ, while Gcn5-dependent H3K36 acetylation is required for HR (Pai *et al*., 2014). As a result, the specific roles of different forms and types of H3K36 modifications in HR or NHEJ repair pathways are still not fully understood.

In this study, we have elucidated the distinct roles of H3K36me2 and H3K36me3 in facilitating specific DSB repair pathways. We showed that two key yeast repair proteins, yKu70 and Rfa1 (a subunit of the RPA complex), are selectively recruited to the damaged chromatin locus by their recognition of H3K36me2 or H3K36me3, respectively. This targeted recruitment subsequently promotes either NHEJ or HR repair efficiency. Moreover, our investigations extended to human homologous proteins, KU70 and RPA1, which bear striking structural similarities to their yeast counterparts. We found that these human proteins also bind to H3K36me2 and H3K36me3 and promotes NHEJ or HR repair efficiency, respectively, suggesting a potential conserved mechanism across species. Overall, our findings unveil that different methylation states on a single histone H3 Lys36 residue can influence the choice of DSB repair pathway, and advance our understanding of the molecular mechanisms of this epigenetic modification in DSB repair.

## Results

### Set2-mediated H3K36 di- and tri-methylation may play different roles in DSB repair

To interrogate whether different forms of H3K36 methylation participate in different DSB repair pathways, a system with Phe/Tyr switch mutations of Set2 was taken advantaged in budding yeast (DiFiore *et al*., 2020, Mei *et al*., 2021). In this assay, H3K36 methylation states of two Set2 mutations, Y149F and Y236F, which uniquely impair di- or tri-methylation of H3K36 (hereafter referred to as Y149F^me3^ and Y236F^me2^, as they exclusively bear H3K36me3 or H3K36me2), and an N-terminal truncation of Set2_1-618_, which impairs both di- and trimethylation of H3K36, were validated by Western blots (Appendix Fig S1A). In addition, the *RAD51* or *yKU70* gene essential for HR or HNEJ repair was further deleted in these strains, allowing for the individual investigation of H3K36me’s role in the NHEJ or HR pathway. Under these scenarios, *rad51Δ* cells or *yku70Δ* cells treated with various recognized DSB-induced agents, including bleomycin, camptothecin (CPT), and etoposide (VP-16), all exhibited severe sensitivity. Moreover, the double deletion of *RAD51 or yKu70* with *SET2* exacerbated the phenotypes, suggesting that Set2 contributes to resistance to DSB (Fig 1 A and B). Interestingly, only add-back of Y236F^me2^, but not Y149F^me3^ and Set2_1-618_, in the HR-deficient *rad51Δset2Δ* strains restored drug resistance nearly to that of wild-type (WT) Set2-expressing cells (Fig 1A). Accordingly, add-back of Y149F^me3^, but not Y236F^me2^ and Set2_1-618_, in the NHEJ-deficient y*ku70Δset2Δ* strains restored drug resistance nearly to that of cells expressing WT Set2 (Fig 1B). Of note, Set2_1-618_, which only retains H3K36me1 could not rescue *SET2* deficiency-caused sensitivity defects, suggesting that H3K36me1, may be dispensable for DSB repair. Altogether, these data suggest that H3K36me2 may function in the NHEJ repair pathway, while H3K36me3 may play an additive role in the HR repair pathway.

**Figure 1.**
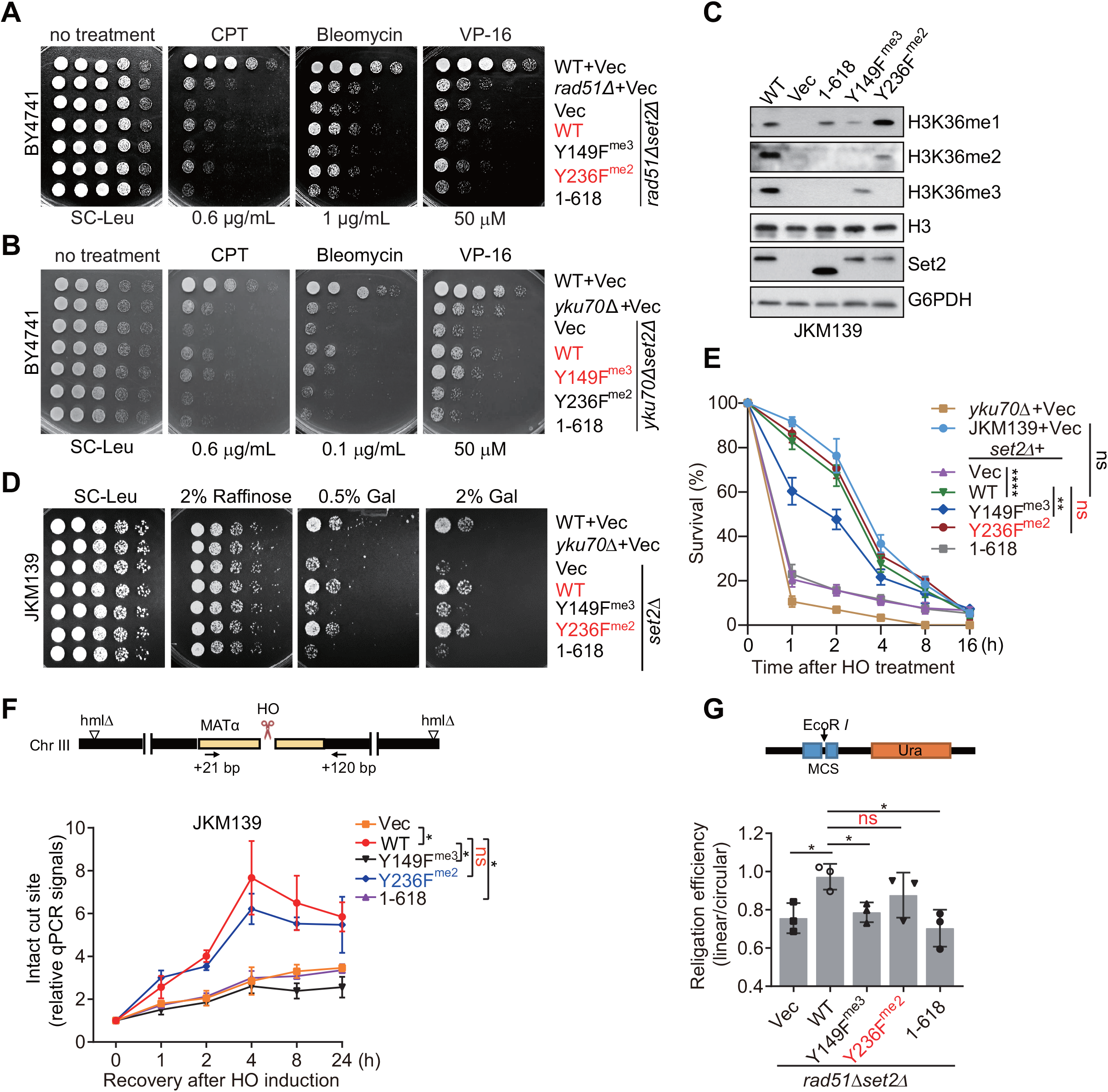
Set2-mediated H3K36me2 and H3K36me3 play distinct roles in DSB repair. A, B. Spotting assays showed the sensitivity of *rad51Δset2Δ* (A) or *yku70Δset2Δ* strains expressing the indicated Set2 mutants in the BY4741 genotype background, when treated with three DSB-inducing drugs. C. Western blot examined the levels of H3K36 methylation in JKM139 strains expressing various Set2 mutants. D. Spotting assays showed the sensitivity of JKM139 strains expressing the indicated Set2 mutants to persistent HO endonuclease-mediated DSB induction by galactose. E. Cell survival populations of the indicated strains were quantified at different time points following HO-induced DSBs. *P*-values represent the significance between different samples at 1- and 2-h HO-induced time points. F. Schematic of HO cut on chromosome III of the haploid yeast to measure its cleavage efficiency using the indicated PCR primers (top panel). Relative qPCR signals at the cutting sites were normalized with the corresponding 0-h time point of each strain, then plotted at the indicated time points upon 1.5-h HO induction following recovery (bottom panel). *P*-values represent the paired difference of each time points between samples. G. Plasmid re-ligation assays were performed in the *rad51Δset2Δ* strains expressing various Set2 mutants. Data information: Data are representative of three independent experiments. Error bars denote the mean ± SD by an unpaired *t*-test. * *P* < 0.05, ** *P* < 0.01, *** *P* < 0.001, ns: not significant.

### H3K36me2 promotes DSB repair via NHEJ pathway

To verify the role of H3K36me2 in NHEJ, the JKM139 strain was utilized (Zhu *et al*., 2008). It has galactose-inducible HO endonuclease, causing site-specific DSBs in chromosome III repaired solely by NHEJ (Appendix Fig S1B). Consistent with the drug sensitivity shown in the BY4741 strain, *set2Δ* exhibited similar sensitivity to galactose-induced DSB as *yku70*Δ cells, rescuable by expressing WT or Y236F^me2^, but not Y149F^me3^ mutant (Figs 1C and D, Appendix Fig S1C). Set2 mutants’ divergent sensitivities weren’t due to carbon source shift, as cells showed comparable growth in raffinose-supplemented media without DSBs (Fig 1D). Alternatively, an ectopic homologous recombination strain tGI354 was also used (Ira & Haber, 2002), where the HO-induced DSB at *MAT* locus of chromosome V is repaired using the homologous *MAT*α-inc sequence at chromosome III as a template (Appendix Fig S1D). In this strain, *set2*Δ cells expressing Y236F^me2^ or Y149F^me3^ exhibited equal DSB sensitivity to WT, confirming irrelevance of carbon source changes for the different phenotypes in JKM139 cells (Appendix Fig S1E).

To demonstrate the DSB repair ability of the Y236F^me2^ mutant via the NHEJ pathway, three different experiments were conducted. First, DSB-induced cell survival assays in JKM139 cells expressing various Set2 constructs were performed ((Zheng *et al*., 2018) details see Materials and Methods). Deletion of *SET2* showed kinetics similar to *yku70Δ* cells, confirming Set2’s role in NHEJ repair. Interestingly, the Y236F^me2^ mutant exhibited kinetics similar to WT Set2, unlike the Y149F^me3^ cells, suggesting that H3K36me2 promotes NHEJ repair (Fig 1E). Secondly, quantitative PCR (qPCR) analysis across the HO recognition site showed that cells expressing WT or Y236F^me2^ Set2 had much more efficient repair than other Set2 mutants after HO-induction recovery. After 4-h recovery, their efficiency was 2-3 times higher than that of the Y149F^me3^ mutant (Fig 1F). Thirdly, a plasmid-based DSB re-ligation assay was performed, in which the linearized p416 plasmid was introduced into *rad51Δset2Δ* cells expressing various Set2 constructs, preventing homologous recombination from occurring (Zheng *et al*., 2018). As expected, Y236F^me2^ showed comparable re-ligation efficiency to WT cells, while Y149F^me3^ displayed 30% lower re-ligation efficiency than *set2Δ* cells (Fig 1G). Importantly, the Y236F^me2^ mutant showed similar chromatin accessibility patterns as WT cells and other Set2 mutants after micrococcal nuclease (MNase) treatment following exposure to bleomycin, as seen through low molecular weight DNA intensity. This indicates that the effect of H3K36me2 on NHEJ repair does not result from altered global chromatin accessibility (Appendix Fig S1F). Moreover, although the Y236F^me2^ mutant displayed considerably delayed cell-cycle progression compared to WT cells after release from G1 arrest, the failure of NHEJ repair in *set2Δ* cells, which have the same delayed cell-cycle progression phenotype, suggested that H3K36me2 bias towards NHEJ is not solely due to G1 arrest (Appendix Fig S1G). Overall, these results illustrate H3K36me2’s important role in promoting NHEJ repair.

### H3K36me3 promotes DSB end resection and HR repair

Next, we explored the potential role of H3K36me3 in the HR repair pathway. To block the NHEJ repair pathway, the *YKU70* gene was deleted in the tGI354 strain, which efficiently utilizes the HR repair machinery. Subsequently, Cells expressing various Set2 constructs were examined for different H3K36 methylation states (Appendix Fig S2A). In this strain, deletion of *YKU70* did not show obvious sensitivity to galactose-induced DSB due to the functioning HR repair pathway. However, further deletion of *SET2* worsened the repair defect (Fig 2A, top panel). Expressing the Y149F^me3^ mutant, but not the Y236F^me2^, in the double knockout cells rescued DSB-sensitive defects, similar to expressing the WT construct, irrespective of different galactose concentrations (Fig 2A, bottom panel). This suggests that H3K36me3, not H3K36me2, may participate in the HR repair process.

**Figure 2.**
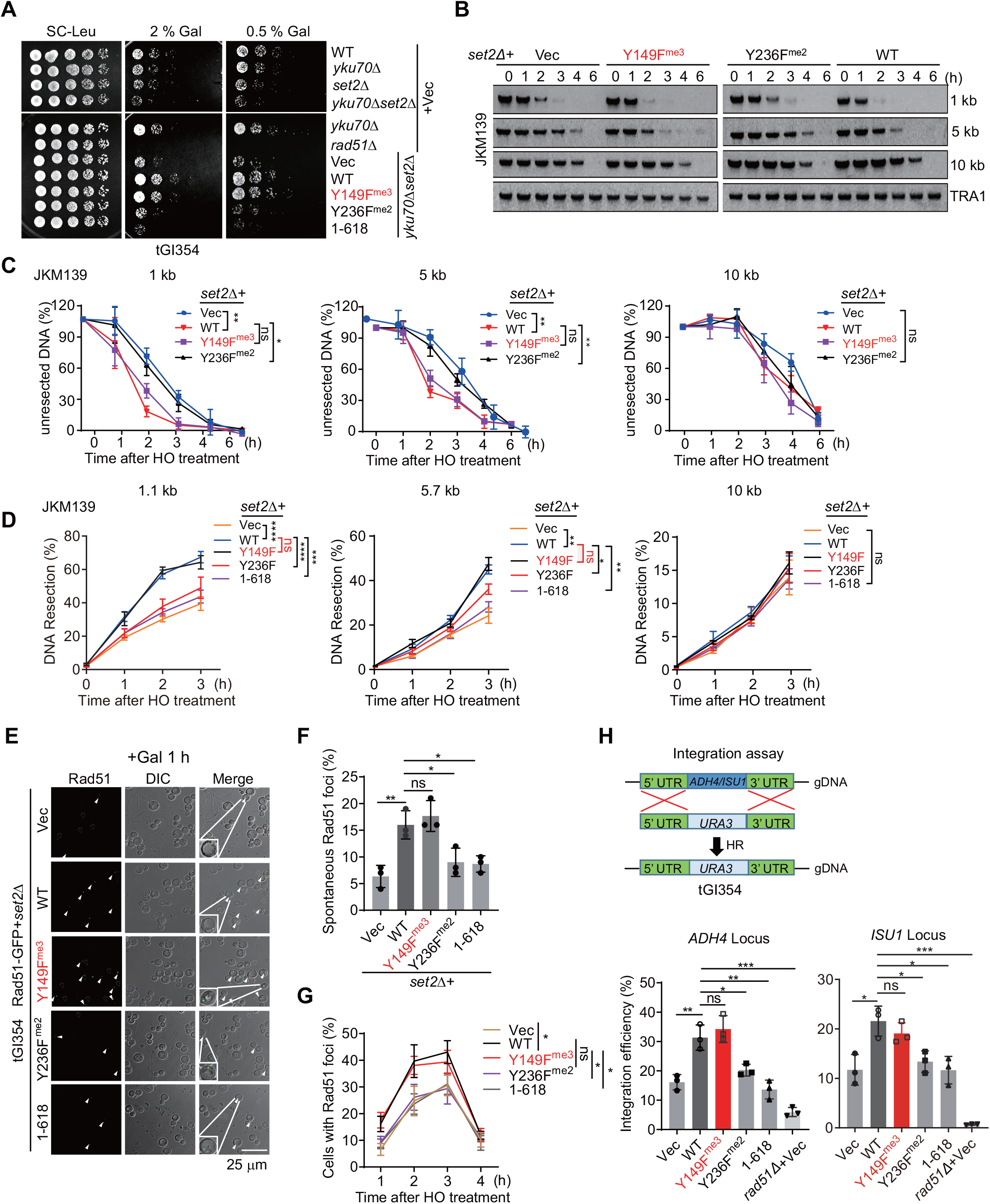
H3K36me3 promotes DSB end resection and HR repair. A. Spotting assays showed the sensitivity of tGI354 strains expressing the indicated Set2 mutants to HO-induction. B, C. Southern blot measured unresected DNAs at the indicated loci away from DSB in *set2Δ* cells expressing various Set2 mutants upon HO-induction of different time points (B) and quantification of DNA resection kinetics of various strains were plotted (C). *TRA1* levels serve as loading controls. D. qPCR analyses were performed to measure DNA resection kinetics upon HO induction. E. Representative microscopy images of *set2Δ* cells expressing various Set2 mutants with Rad51-GFP upon 1-h HO-induction. Rad51-GFP foci are indicated by arrows. Enlarged cells with GFP signal are shown. DIC: differential interference contrast. F. Cell populations with Rad51-GFP foci shown in (E) were quantified. G. Kinetics of cell population with Gad51-GFP foci upon HO-induction for the indicated times in various Set2 mutants were plotted. H. Schematic of integration rate assay by HR to replace *ADH4* or *ISU1* gene with URA3 gene (top panel), and gene integration efficiency in the indicated strains was measured with the colonies ratio (bottom panel). Data information: Data are representative of three independent experiments. Error bars denote the mean ± SD with *P*-value by an unpaired *t*-test. * *P* < 0.05, ** *P* < 0.01, *** *P* < 0.001, ns: not significant.

Next, we examined the role of H3K36 methylation in DSB end resection using JKM139 strain, which cannot repair DSBs by HR. Resection kinetics at the site-specific DSB were monitored using Southern blot or qPCR assays. In qPCR assay, the 5’ strand degradation at DSB ends prevents the restriction enzyme from cleaving single strand DNA (ssDNA), enabling PCR amplification of the remaining DNA fraction (Appendix Fig S2B). The intensity of the bands in Southern blot hybridization indicates unresected DNA fragments (Fig 2B). Y149F^me3^ cells exhibited similar end resection kinetics at both the proximal ends and 10 kb distal ends compared to the WT cells. In contrast, Y236F^me2^ cells exhibited relatively poor resection at the proximal ∼ 1 or 5 kb ends, but no difference at 10 kb distal ends within 3-h HO-induction (Figs 2C and D). The defect of Y236F^me2^ mutant in DSB end resection did not arise from delayed checkpoint activation, as reflected by prompt γH2A hyperphosphorylation, which was similar to the WT cells (Appendix Fig S2C). The defect of Y236F^me2^ in DSB end resection is not due to dramatic changes in cell cycle progression, whether before or after persistent galactose induction (Appendix Fig S2D). Thus, we conclude that H3K36me3, but not H3K36me2, facilitates DSB end resection.

Accumulation of Rad51 foci upon DSB induction represents HR repair progression (Miyazaki *et al*., 2004, Waterman *et al*., 2019). We monitored GFP signals in various Set2 strains expressing Rad51-GFP upon persistent HO-induction by fluorescent microscopy. After 1-h of HO-induction, < 8% cells displayed GFP foci in *set2Δ* or Y236F^me2^ cells. while over 15% of cells exhibited GFP foci in both WT and Y149F^me3^ cells, indicating a faster response to HR repair compared to Set2-deficient cells (Figs 2 E and F). The percentage of cells with GFP foci increased to 40% after 2-h of HO-induction but declined to 12% after 4-h of continuous galactose culture. However, Y149F^me3^ cells remained at a similar HR repair efficiency as WT cell at the 2-h duration (Appendix Figs 2E and F). The divergence between WT and Set2 mutants decreased gradually at the 4-h duration, presumably because the HR-fixed locus was no longer cut under continuous HO-induction (Fig 2G). Moreover, the integration assay was employed to measure HR repair efficiency by detecting the integration rate of the *URA3* gene at the *ADH4* or *ISU1* locus via HR. As expected, Y149F^me3^ exhibited a comparable integration efficiency to WT cells, whereas H3K36me3-deficient mutants did not (Fig 2H). These data demonstrate that H3K36me3, but not H3K36me2, promotes DSB end resection and HR repair.

### yKu70 uniquely interacts with H3K36me2 *in vitro*

Based on above results, we propose that H3K36me2 and H3K36me3 have distinct roles in response to different DSB repair signaling. To this end, we reviewed known critical regulators associated with DDR and DNA repair, and identified several abundant cellular proteins (Appendix Fig S3A). We generated various integrated epitope-tagged strains, and surveyed the interactions of these proteins with histone H3. Interestingly, we found that yKu70, Rfa1, and Xrs2 proteins were able to co-immunoprecipitate (co-IP) with histone H3 (Appendix Figs S3B and C). Reciprocal co-IP experiments confirmed that H3K36me2 or H3K36me3 binds with yKu70 and Rfa1, respectively (Appendix Figs S3D and E). Surprisingly, yKu70 specifically interacts with H3K36me2, but not H3K36me3, and their interaction was enhanced upon HO-induction or bleomycin treatment (Fig 3A-C).

**Figure 3.**
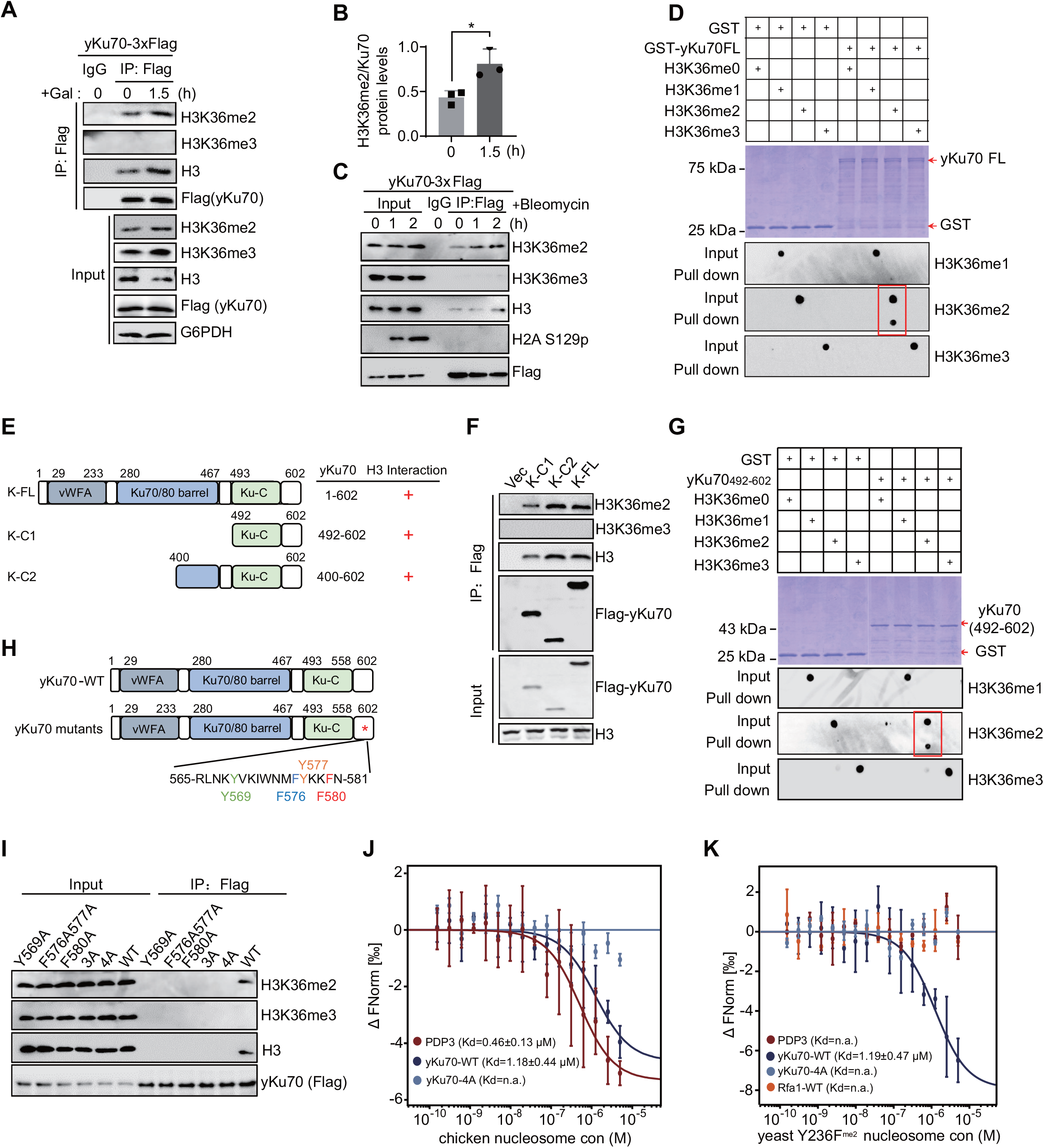
Identification of the binding segment of yKu70 towards H3K36me2 *in vitro*. A, B. Co-IP experiments showed an enhanced interaction of yKu70 with H3K36me2, but not H3K36me3, upon HO-induction (A). Relative H3K36me2 levels bound with yKu70 were quantified (B). C. Co-IP experiments showed that the interaction of yKu70 and H3K36me2 was gradually increased with bleomycin treatment. D. *In vitro* GST pull-down assays to examine the interaction of full-length GST-yKu70 with the indicated methyl-H3K36 peptides by dot blotting. E, F, H, I. Schematic of the full-length or truncated yKu70 proteins used for co-IP assay (E and H), and the interactions of various yKu70 truncations or mutations with H3K36me2 in BY4741 cells were examined (F and I). G. *In vitro* GST pull-down assays were performed to examine the interaction of GST-yKu70_492-602_ with the indicated methyl-H3K36 peptides by dot blotting. GST protein served as a negative control. J, K. Normalized MST binding curves of native chicken erythrocyte nucleosomes (J) or yeast Y236F^me2^ nucleosomes (K) with the WT or 4A mutant of yKu70. The K_d_ values are present, and n.a. represents not determined. Data information: Data are representative of three (B) independent experiments. Error bars denote the mean ± SD by an unpaired *t*-test. * *P* < 0.05.

To verify direct binding between yKu70 and K36me-decorated histone H3, *in vitro* GST pull-down assays were conducted using recombinant GST or GST-yKu70 incubated with various synthetic H3K36 peptides. Consistently, GST-yKu70 specifically bound to the H3K36me2 peptide, indicative of its specificity (Fig 3D). We also observed that yKu80 barely bound to H3K36me2 (Appendix Fig S4A).

We further sought to determine the precise H3K36me2-associated region(s) on the yKu70 protein. A series of truncated yKu70 fragments were generated, and their binding abilities were systematically examined. We found that the C-terminus of yKu70 is important for binding with H3K36me2 (Appendix Figs S4B and C, Figs 3E and F). GST pull-down and peptide pull-down assays confirmed the requirement of the C-terminal yKu70 fragments in the interaction of H3K36me2 (Fig 3G, Appendix Fig S4D). Additionally, we further confirmed that the region containing amino acids_559-580_ in yKu70 is critical for interacting with histone H3 (Appendix Fig S4E-H).

An aromatic cage, typically formed by Phe/Tyr/Trp residues, is critical for recognizing methylated-lysine (Qin & Min, 2014). To investigated the key residues within this fragment required for H3K36me2 recognition, structure docking suggested that an aromatic cluster composed of four residues on yKu70 (Appendix Fig S4I). Indeed, mutating any of residues (Y569A, F576AY577A, and F580A) or a combination of F576/Y577/F580 to alanine (3A mutant), or mutating all four residues (4A mutant), resulted in the loss of binding ability towards histone H3 or H3K36me2 (Fig 3H and I). To determine if yKu70 directly binds H3K36me2-decorated nucleosomes, *in vitro* quantitative microscale thermophoresis (MST) binding assay was used. Ideally, reconstituted H3K36me2-modified nucleosome is the best material for this assay. Unfortunately, there is no commercial product available, and currently it is technically difficult to generate this homogenous H3K36me2-nucleosome on the bench. Therefore, native purified nucleosomes were used in this assay, in which various histone modifications are preserved. However, if loss of H3K36me2 marks itself is sufficient to prevent the binding of yKu70 and H3K36me2 nucleosome, one would predict that H3K36me2 is essential for this association, regardless of other existed histone modifications. Indeed, MST experiments showed that the WT yKu70 displayed a comparable binding affinity towards both purified native chicken erythrocyte nucleosomes and yeast Y236F^me2^ nucleosomes, in contrast to undetectable binding affinity displayed by the 4A mutant or WT Rfa1 protein (Kd ≈ 1.2 μM in WT vs. not detected in 4A mutant, Figs 3J and K, Appendix Fig S5A and B). Notably, the PDP3 protein contains a PWWP domain that tightly bind H3K36me3 *in vitro* was used as a control (Gilbert *et al*., 2014). These results indicate that yKu70 directly binds to H3K36me2 mainly via four essential aromatic amino acid residues.

### H3K36me2 recruits yKu70 at DSB to facilitate DNA repair through NHEJ

As the yKu complex can be recruited to damaged chromatin during DNA damage (Fnu *et al*., 2011), we performed chromatin immunoprecipitation (ChIP)-qPCR analysis to assess the ability of H3K36me2 in yKu70 recruitment *in vivo*. We observed that yKu70 was mainly enriched at the proximal 1-kb region away from the HO-cut site. Interestingly, H3K36me2 and Set2 levels also increased at the 1-kb locus upon HO-induction (Figs 4A and B). These results were supported by the occupancy of DNA ligase IV (Dnl4/Lif1), required for NHEJ repair to ligate the break ends (Zhang *et al*., 2007), across the DSB genome locus and peaking at the same proximal 1-kb locus upon longer HO-induction (Fig 4C). Concordantly, ChIP-qPCR assay showed that the 4A mutant abated yKu70 enrichment at the 1-kb locus of DSB but barely affected H3K36me2 occupancy at the same loci (Fig 4D). These data indicate that yKu70 enrichment occurs downstream of H3K36me2 accumulation upon DNA damage. Supportively, the binding-deficient mutants of yKu70, especially 4A mutant, exhibited severe sensitivity to galactose-induced DSB or various drugs, compared to cells expressing WT yKu70 (Figs 4E and 4F). Thus, these data suggest that H3K36me2 facilitates the recruitment of yKu70 to the proximal DSB locus, facilitating NHEJ repair.

**Figure 4.**
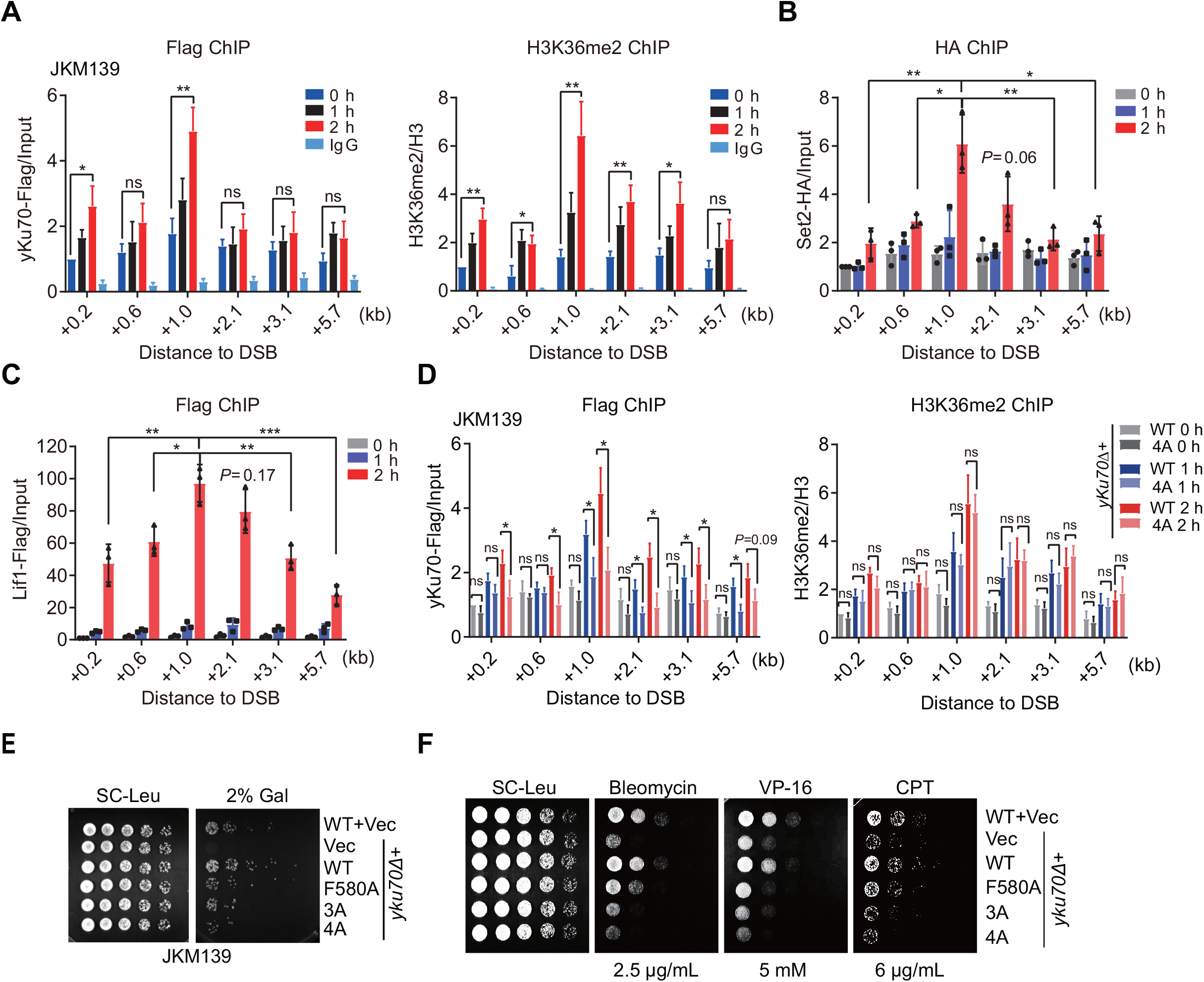
Recruitment of yKu70 by H3K36me2 at the proximal locus of DSB to promote NHEJ repair. A. ChIP-qPCR showed the enrichment of yKu70-Flag or H3K36me2 at different loci away from the DSB site with the indicated HO-induced times. B, C. ChIP-qPCR showed the enrichment of Set2-HA (B) or Lif1-Flag (C) at different loci away from the DSB site with the indicated HO-induced times. D. ChIP-qPCR showed the enrichment of yKu70-Flag or H3K36me2 in strains expressing WT or 4A mutant at different loci upon HO-induction at different times. E. Spotting assay showed the sensitivity of JKM139 strains expressing the indicated yKu70 mutants upon galactose induction. F Spotting assay showed the sensitivity of JKM139 strains expressing the indicated yKu70 mutants upon treatment with three DSB-induced drugs. Data information: Data are representative of three independent experiments. Error bars denote the mean ± SD by an unpaired *t*-test. * *P* < 0.05, ** *P* < 0.01, ns, not significant.

### Rfa1 specifically interacts with H3K36me3 *in vitro*

Rfa1 exclusively interacts with H3K36me3 chromatin but not H3K36me2 (Appendix Fig S3C). We propose that H3K36me3 may directly interact with RPA. We confirmed that the interaction of Rfa1 with H3K36me3 is enhanced upon DNA damage induction (Figs 5A-C). GST pull-down assays also showed that only the Rfa1 subunit directly binds synthetic H3K36me3 peptide, not Rfa2 or Rfa3 subunits (Fig 5D and Appendix Fig S6A).

**Figure 5.**
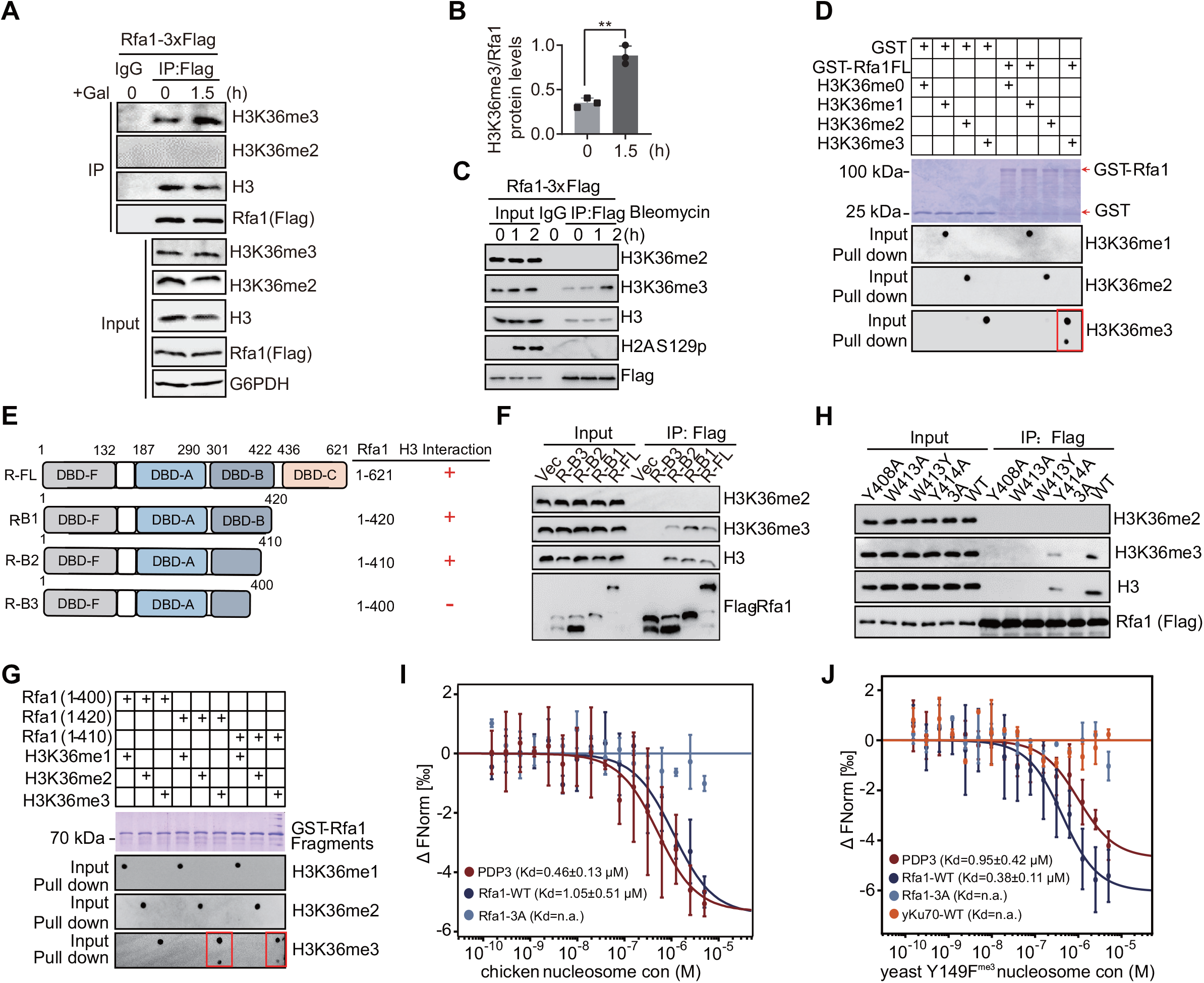
Identification of the binding segment of Rfa1 towards H3K36me3 *in vitro*. A, B. Co-IP showed an enhanced interaction of Rfa1 with H3K36me3, but not H3K36me2, upon HO-induction (A). Relative H3K36me3 levels bound with Rfa1 were quantified (B). C. Co-IP showed that the interaction of Rfa1 and H3K36me3 was gradually increased with bleomycin treatment. D, G. *In vitro* GST pull-down assays to examine the interaction of full-length GST-Rfa1 (D) or truncated forms (G) with the indicated methyl-H3K36 peptides by dot blotting. E, F. Scheme showing full-length or truncated Rfa1 proteins used for co-IP assay (E), and their interactions with H3K36me3 in BY4741 cells were examined (F). H. Co-IP showed the interacting abilities of various Rfa1 mutant proteins towards H3K36me3. I, J. Normalized MST binding curves of native chicken erythrocyte nucleosomes (I) or yeast Y149F^me3^ nucleosomes (J) with the WT or 3A mutant of Rfa1. The K_d_ values are present, and n.a. represents not determined. Data information: Data are representative of three (B) independent experiments. Error bars denote the mean ± SD by an unpaired *t*-test. ** *P* < 0.01.

Through a series of experiments using truncated Rfa1 proteins, we aimed to identify the precise region(s) responsible for binding to H3K36me3. Our results revealed that the DBD-B (DNA-binding domain-B) domain of Rfa1 plays a crucial role in this interaction (Appendix Fig S6B-E). Removal of this domain completely abolished the binding of Rfa1 to H3K36me3 (Appendix Figs S6F and G). Further narrowing down the interacting region, we pinpointed that the residues 400-410 within Rfa1 are essential for this interaction (Figs 5E and F). *In vitro* GST pull-down assay also confirmed that the absence of these amino acids (401– 409) impaired the association of Rfa1 with synthetic H3K36me peptide (Fig 5G).

Three aromatic residues of Rfa1 in recognizing H3K36me3 were identified through structural docking the Rfa1 fragment_336–426_ with the H3K36me3 peptide using Autodock software, revealing three aromatic residues (Appendix Fig S6H). Co-IP experiments showed that all three mutants attenuated Rfa1 binding with histone H3 and H3K36me3, but neither interacting with H3K36me2 (Fig 5H). Peptide pull-down assays confirmed specific binding of the H3K36me3 peptide to WT Rfa1, not the aromatic mutant (Appendix Fig S6I). Likewise, native nucleosomes were used to determine the direct interaction between Rfa1 and H3K36me3-decorated nucleosomes. MST experiments revealed a strong binding affinity of WT Rfa1 with native chicken erythrocyte nucleosomes and yeast Y149F^me3^ nucleosomes, while the 3A mutant and WT yKu70 protein showed undetectable binding (Kd ≈ 1.0 μM in WT vs. not detected in 3A mutant, Figs 5I and J; Appendix Figs S5A and C). The results indicate that Rfa1 directly interacts with H3K36me3 mainly via three aromatic amino acid residues.

### H3K36me3 recruits RPA to the nearby locus of DSB and facilitates HR repair

Next, we sought to examine whether RPA could be recruited to the DSB by H3K36me3 *in vivo*. Therefore, ChIP-qPCR was performed and we unexpectedly detected a peak signal of Rfa1 at the 3.0-kb locus away from the DSB, coinciding with H3K36me3 and Set2 peaks, implying a potential association of H3K36me3 and RPA in this region (Figs 6A and B). The Rad51 protein, which promotes the formation of homologous exchange filament, was also found to occupy the proximal DSB genome locus and peaked at the similar location as Rfa1 upon HO-induction (Fig 6C). Concordantly, ChIP-qPCR assay indicated that the 3A mutant disassociated Rfa1 from the 0.2-kb and 3.0-kb loci away from the DSB, but did not influence H3K36me3 enrichment at the same loci, suggesting downstream recruitment of RPA to H3K36me3 accumulation (Fig 6D). Spotting assays further confirmed that the H3K36me3 binding-deficient Rfa1 mutants exhibited severe sensitivity towards galactose-induced DSB or various drugs, compared with cells expressing WT RPA1 (Figs 6E and F). It is worthy to point out that DNA resection from the DSB site does not proceed equally in each cell, which may result in a subpopulation of cells being unresected at around 3-kb locus away from DSB even after 2-h of HO-induction. This notion was supported by the results in DSB end resection assays shown above (Fig 2B-D). In the end, preserved chromatin with certain signatures nearby DSBs, but not distal regions of DSBs, may serve as a platform to recruit the RPA complex and other damage-response factors to assist DNA repair. In summary, these data suggest that H3K36me3 is critical for recruiting subpopulation of RPA to the nearby chromatin of DSB.

**Figure 6.**
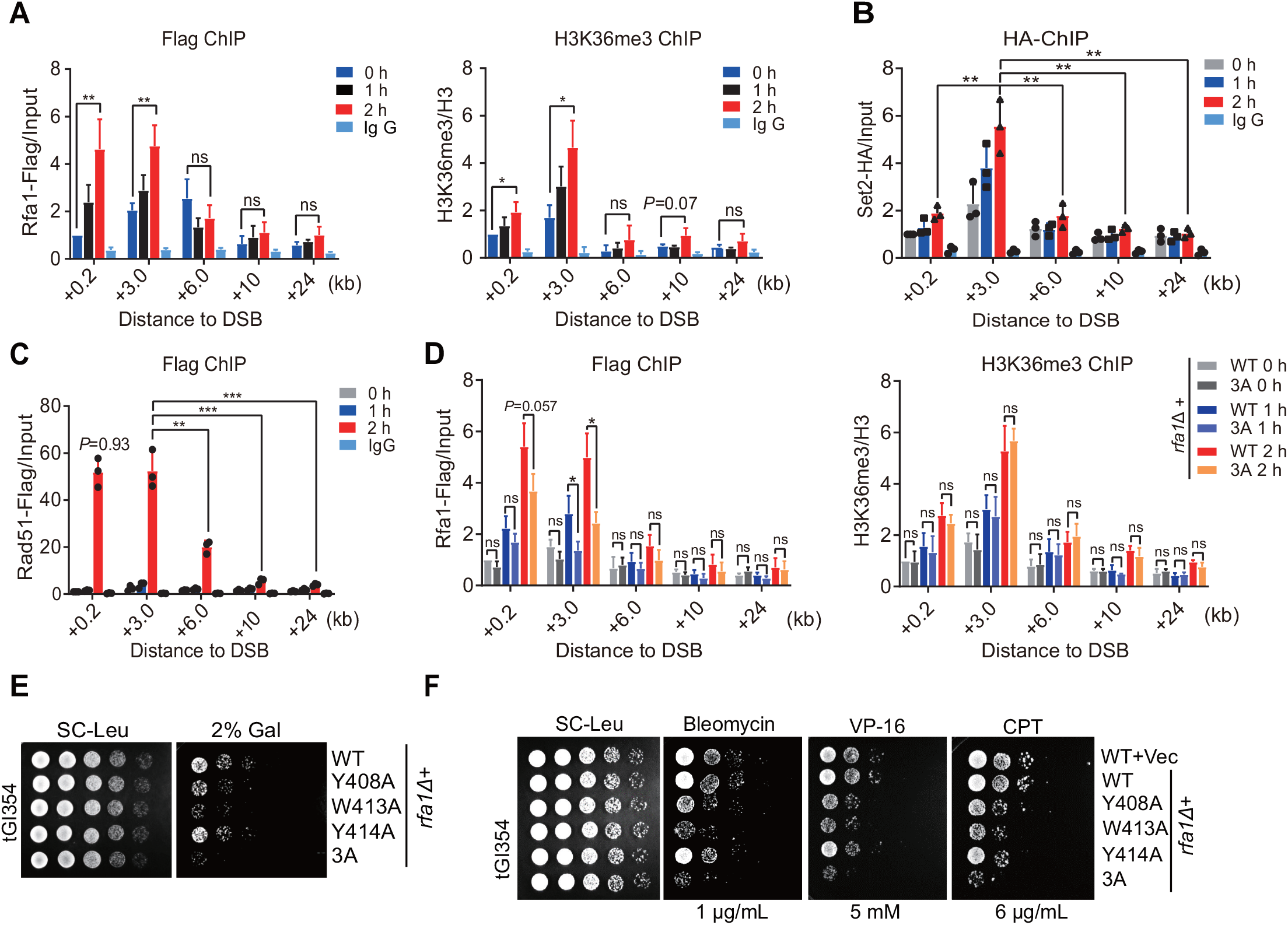
Recruitment of Rfa1 by H3K36me3 at the locus nearby DSB to facilitate HR repair. A. ChIP-qPCR showed the enrichment of Rfa1 or H3K36me3 at various loci away from DSB at the indicated time points. B, C. ChIP-qPCR showed the enrichment of Set2-HA (B) or Rad51-Flag (C) at different loci away from the DSB site with the indicated HO-induced times. IgG ChIP samples served as negative controls. D. ChIP-qPCR showed the enrichment of Rfa1-Flag or H3K36me3 in strains expressing the WT or 3A mutant strain at different loci upon HO-induction at different times. E. Spotting assays showed the sensitivity of various tGI354 strains expressing the indicated Rfa1 mutants by HO-induction. F. Spotting assays showed the sensitivity of various tGI354 strains expressing the indicated Rfa1 mutants treated with three DSB-induced drugs. Data information: Data are representative of three independent experiments. Error bars denote the mean ± SD by an unpaired *t*-test. * *P* < 0.05, ** *P* < 0.01, ns, not significant.

### The differing genomic distributions of H3K36 methylation marks controls the recruitment of yKu70 or Rfa1 upon DSB induction

H3K36me2 and H3K36me3 co-occupy at active transcribed gene bodies. However, recent studies indicate that, in mammalian cells, H3K36me3 exhibits characteristic enrichment within gene bodies, while H3K36me2 displays a more diffuse distribution across both genic and intergenic regions, contributing to shaping the DNA methylation landscape (Weinberg *et al*., 2019, Yano *et al*., 2022). Consistent with this, our analysis of the public GSE116646 dataset revealed that H3K36me3 is predominantly enriched at gene bodies, while H3K36me2 is found in both genic and intergenic regions (Gopalakrishnan *et al*., 2019). This suggest that H3K36me2 and H3K36me3 may exclusively recruit yKu70 or Rfa1 at different genomic regions. To test this hypothesis, we induced DSBs in yeast cells using bleomycin and performed ChIP-qPCR for H3K36me2/me3, yKu70 and Rfa1 at several gene bodies with higher H3K36me3 enrichment and adjacent intergenic regions decorated with H3K36me2. Interestingly, in response to DSB stress, we observed that H3K36me2 levels were gradually increased at intergenic regions, but not at transcribed gene bodies, with a similar trend seen for yKu70 enrichment, regardless of the genome loci. On the other hand, H3K36me3 levels specifically increased at transcribed gene bodies but not at intergenic regions, similar to the pattern observed for Rfa1 enrichment (Appendix Figs S7A and B). The successful induction of DSBs was validated by examining γH2A.X levels (Appendix Fig S7C). Based on these findings, we propose that transcriptionally active genomic regions may preferentially utilize error-free HR repair to maintain genome fidelity, while intergenic regions may prefer faster NHEJ repair way to promote cell proliferation. This highlights the distinct roles of H3K36me2 and H3K36me3 in guiding the recruitment of specific DSB-related effectors for regulating DSB repair pathways.

### Human KU70 and RPA1 exclusively bind with H3K36me2 and H3K36me3 in a conserved manner

The primary sequences and tertiary structures of yeast Rfa1 and yKu70 are conserved in higher eukaryotes. Alignment of yKu70 with homologous proteins revealed conservation of F580 (Appendix Fig S8A). Crystal structures of yKu70 and human KU70 showed domain conservation, except for a separate module in KU70, as highlighted (Appendix Fig S8B). Similarly, alignment of yeast Rfa1 with orthologs from different species revealed high conservation of Y408, W413, and Y414 (Appendix Fig S8C). Crystal structures of Rfa1 and RPA1 exhibited significant similarity, including methyl-lysine binding regions (Appendix Fig S8D).

Based on these similarities, we investigated whether human RPA1 and KU70 play similar roles in mammalian cells. As anticipated, our results demonstrated that RPA1 specifically binds to H3K36me3 peptides, while KU70 interacts exclusively with H3K36me2 peptides (Fig 7A). Reciprocal *in vitro* peptide pull-down assays further confirmed these specific interactions (Fig 7B). In cells, Flag-tagged RPA1 and KU70 exclusively bound with H3K36me2 or H3K36me3, respectively, confirming the functional conservation (Fig 7C). Co-IP experiments with conserved mutants of KU70 or RPA1 validated the significance of the aromatic cage-like centers, showing reduced binding to H3K36me2 or H3K36me3, respectively (Figs 7D and E). These data support the conservation action of yeast proteins and their human orthologs in recognizing H3K36me2 and H3K36me3 marks.

**Figure 7.**
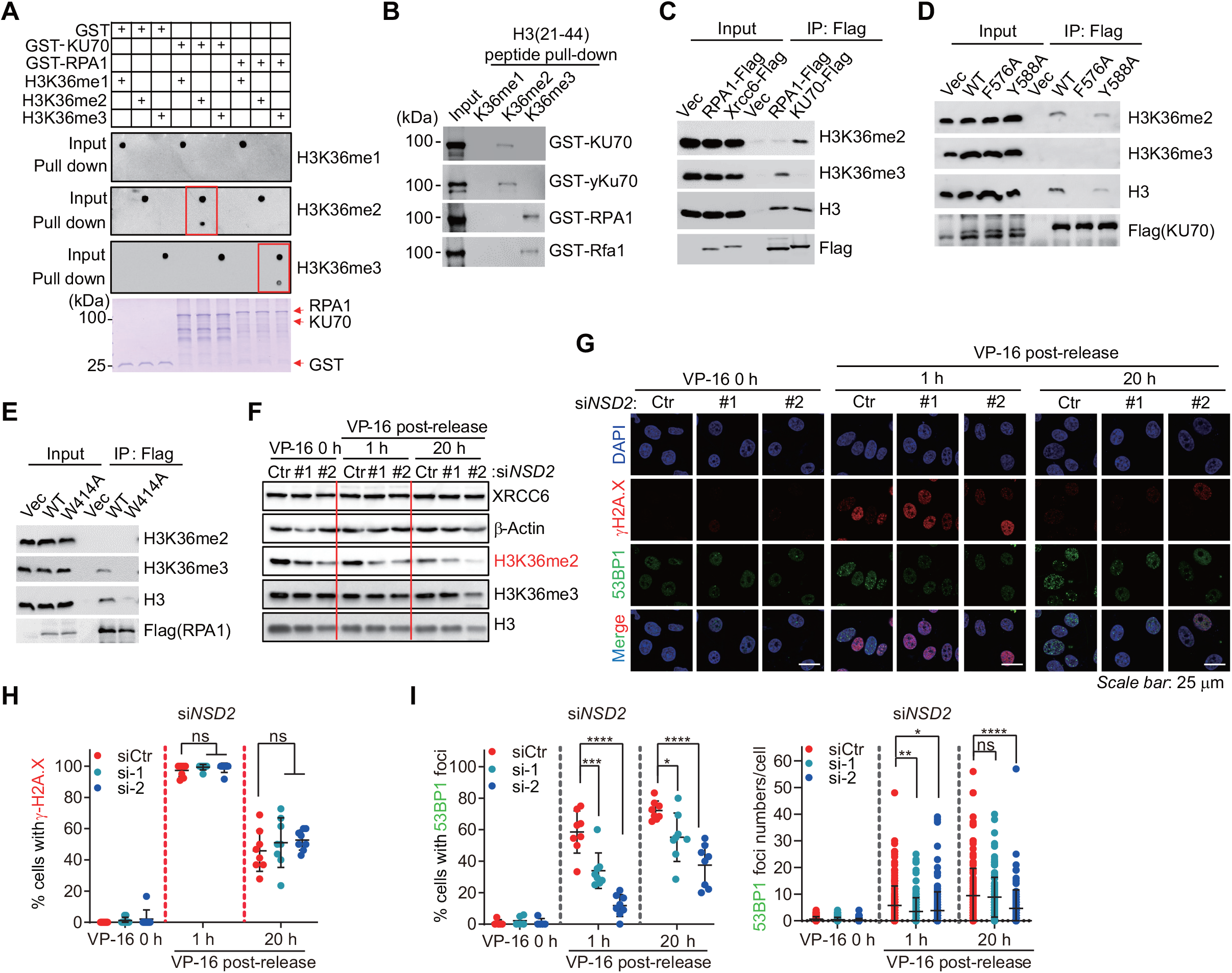
H3K36me2 interacts with human KU70 and promotes 53BP1 accumulation at DSBs. A. *In vitro* GST pull-down assays to examine the interaction of full-length GST-KU70 or GST-RPA1 with the indicated methyl-H3K36 peptides by dot blotting. B. *In vitro* peptide pull-down assays showed that different H3K36 methyl peptide binds to GST-KU70 or GST-RPA1 protein, respectively. C-E. Co-IP showed KU70 or RPA1 binds H3K36me2 or H3K36me3 in HEK293T cells, respectively (C), and various mutant of KU70 (D) or mutants of RPA1 (E) disrupt their binding to methyl-H3K36 nucleosomes, respectively. F. U-2OS cells transfected with si*RNA* Control (Ctr) or two si*RNA* oligos targeting *NSD2* were treated with VP-16 (50 μM) for 1 h, and then released into normal medium for the indicated times. Knockdown efficiency of *NSD2* was examined by immunoblotting with H3K36me2. G-I. Representative Immunostaining images with γH2A.X and 53BP1 in cells with the indicated treatment shown in (F) were presented (G). γH2A.X-positive cells (H) and 53BP-positive cells and foci numbers per cell (l) were quantified. Data information: Data are representative of cells from 8 different areas (H and I, left panel) and foci numbers from 85 cells (I, right panel) using ImageJ. *Scale bar*, 25 μm (G). Error bars denote the means ± SD with *P*-values determined by an unpaired *t*-test. * *P* < 0.05, ** *P* < 0.01, ** \**P* < 0.001, **** *P* < 0.0001, ns, not significant.

### NSD2-mediated H3K36me2 is critical for the recruitment of 53BP1 at DSBs

In mammalian cells, several studies have shown that the competition between BRCA1/CtIP and 53BP1/Shieldin determines the choice between NHEJ or HR pathways for DSB repair (Bunting *et al*., 2010, Escribano-Diaz *et al*., 2013, Findlay *et al*., 2018). To further validate the roles of H3K36me2 and H3K36me3 in human DSB repair, the methyltransferases NSD2 and SETD2 were depleted, which have been proven to be mainly responsible for H3K36me2 and H3K36me3, respectively (Hu *et al*., 2010, Sengupta *et al*., 2021). Their impacts on different DSB repair pathways were investigated individually. Knockdown of *NSD2* reduced H3K36me2 levels, but not H3K36me3 levels (Fig 7F). Upon treatment with the DSB-inducing reagent VP-16, equivalently enhanced γH2A.X levels were observed in both WT and si*NSD2* cells, suggesting NSD2 is not necessary for response of DSB damage (Figs 7G and H). Interestingly, the formation of 53BP1 foci at DSBs was significantly decreased in si*NSD2* cells after 1 hour of VP-16 treatment (Figs 7G and I). By contrast, depletion of the H3K36me3 methyltransferase SETD2 did not notably affect the number of 53BP1 foci at DSBs with VP-16 treatment (Appendix Figs S9A and B). Supportively, previous studies have correlated H3K36me2 to NHEJ by interrupting potential H3K36me2 methyltransferases, SETMAR or NSD2/MMSET, resulting in limited recruitment of NHEJ proteins and less repair efficiency in mammalian cells (de Krijger *et al*., 2020, Fnu *et al*., 2011, Kaur *et al*., 2020). Based on these findings, we propose that H3K36me2 marks promotes the accumulation of 53BP1 and NHEJ repair at DSBs in human cells.

### SETD2-mediated H3K36me3 is critical for RPA recruitment at DSBs

Furthermore, we investigated the impact of H3K36me3 levels on DSB repair pathways under VP-16 stress. Knockdown of *SETD2* significantly reduced H3K36me3, but not H3K36me2 (Fig 8A). Immunostaining showed that the reduction of H3K36me3 did not affect γH2A.X signals, but remarkably attenuated RPA2 accumulation at DSBs after 1 hour of post-release from VP-16 treatment (Fig 8B-D). These observations are in agreement with previous studies showing that SETD2-mediated H3K36me3 is necessary for HR repair of DSBs induced by IR or I-Sec *I* expression (Carvalho *et al*., 2014, Pfister *et al*., 2014). Importantly, the effect of H3K36me3 on RPA recruitment is specific, as knockdown of *NSD2* did not decrease, or even slightly increased, RPA2 foci numbers at DSBs (Appendix Figs S10A and B). This indicates that impairment of the HR pathway may modulate H3K36me3, potentially favoring the more efficient utilization of the H3K36me2-mediated NHEJ pathway. Overall, our data establish a conserved role of RPA1 and KU70 in the respective recognition of H3K36me3 or H3K36me2 to promote HR or NHEJ repair efficiency.

**Figure 8.**
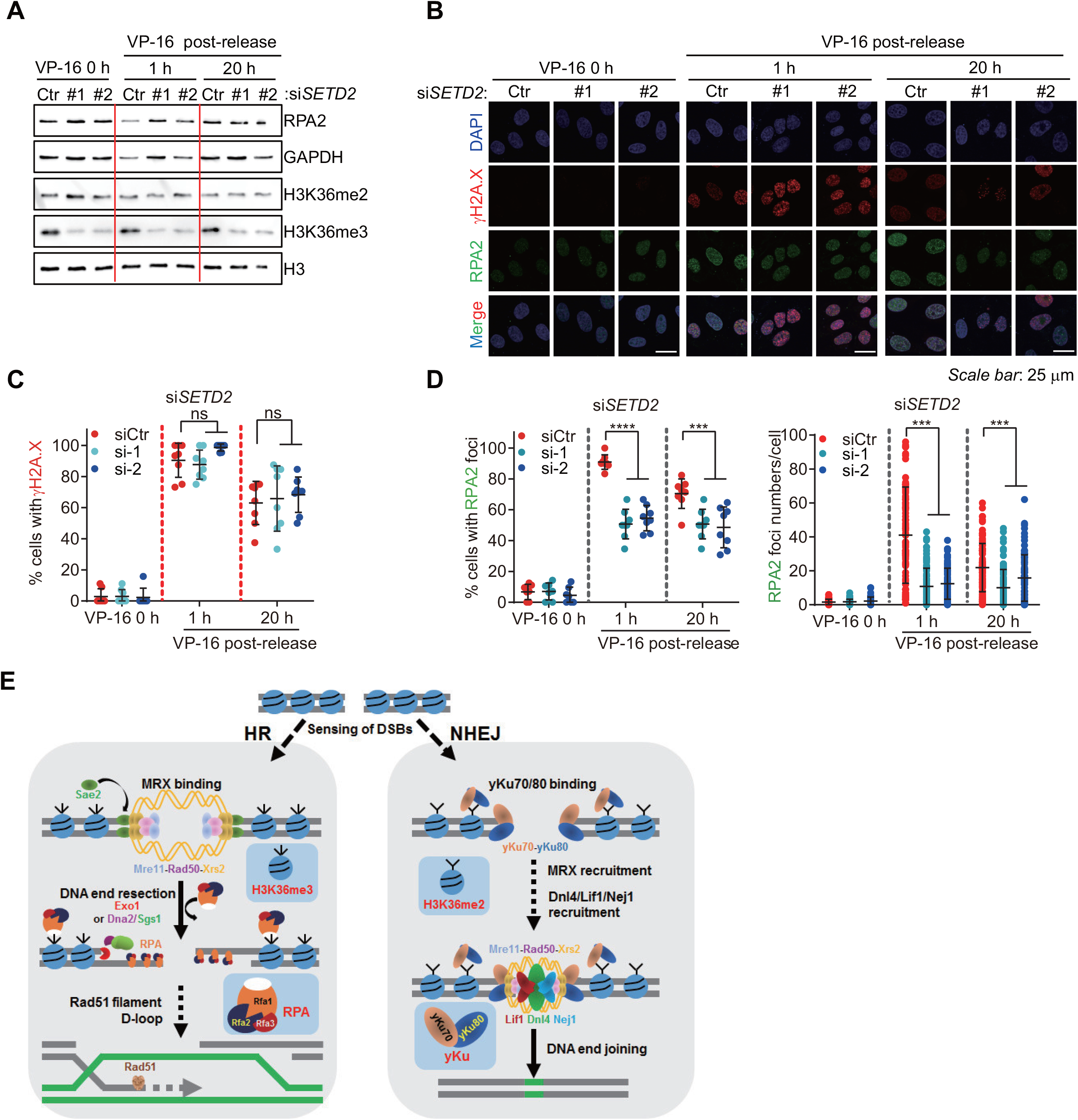
H3K36me3 promotes RPA recruitment at DSBs. A. U-2OS cells transfected with si*RNA* Control (Ctr) or two si*RNA* oligos targeting *SETD2* were treated with VP-16 (50 μM) for 1 h, and then released into normal medium for the indicated times. Knockdown efficiency of *SETD2* was examined by immunoblotting with H3K36me3. B-D. Representative Immunostaining images with γH2A.X and RPA2 in cells with the indicated treatment shown in (A) were presented (B). γH2A.X-positive cells (C) and RPA2-positive cells and foci numbers per cell (D) were quantified. E A working model of H3K36me2 or H3K36me3 recruiting yeast yKu70 (human KU70) or yeast Rfa1 (human RPA1) at DSB sites to promote NHEJ or HR repair pathways, respectively. Data information: Data are representative of cells from > 7 different areas (C and D, left panel) and foci numbers from > 60 cells (D, right panel) using ImageJ. *Scale bar*, 25 μm (B). Error bars denote the means ± SD with *P*-values determined by an unpaired *t*-test. * *P* < 0.05, ** *P* < 0.01, ** \**P* < 0.001, **** *P* < 0.0001, ns, not significant.

## Discussion

Although Set2/SETD2 and its-mediated H3K36 methylation have been implicated in DNA damage and repair, the concomitance of all three H3K36 methylation states in living cells hinders the disclosure of their unique functions in diverse DSB repair pathways. Hence, using budding yeast, we clearly demonstrate that H3K36me2 and H3K36me3 have distinct roles in DSB repair. During NHEJ process, H3K36me2 is enriched at the proximity of DSB, thereby interacting with yKu70 and leading to the chromatin recruitment of the Ku complex to facilitate DSB repair. In contrast, during HR repair, the essential component Rfa1 is not only targeted to ssDNA at the proximal DSB site to maintain genome fidelity but is also enriched at the nearby chromatin locus of the DSB through H3K36me3 recruitment to facilitate HR repair (Fig 8E). More strikingly, we have identified that the human counterparts, RPA1 and KU70, physically interact with H3K36me2 or H3K36me3, respectively. This interaction may likewise function in distinct DSB repair pathways, highlighting the conservation of H3K36me-dependent mechanisms in DNA repair.

### The natural genomic distributions of different H3K36 marks may influence the selection of DSB repair pathway

The chromatin structure decorated with H3K36me plays a role in balancing NHEJ and HR, thereby safeguarding genome stability and organism fitness (Carvalho *et al*., 2014, de Krijger *et al*., 2020, Fnu *et al*., 2011, Jha & Strahl, 2014, Pai *et al*., 2014, Pfister *et al*., 2014). H3K36me2 marks the intergenic regions and recruits the DNA methyltransferase DNMT3A, shaping the DNA methylation landscape. In contrast, H3K36me3 is specifically enriched within active gene bodies (Shirane *et al*., 2020, Weinberg *et al*., 2019). Actually, the mammalian genome comprises > 50% intergenic regions, whereas only 30% of the budding yeast genome are intergenic regions (Lu & Lin, 2019). Thus, it is interpretable that HR repair serves as the primary DSB repair pathway in *S. cerevisiae*, whereas NHEJ is the dominant repair mechanism in higher eukaryotes (Li *et al*., 2019, Scully *et al*., 2019). Thus, we hypothesize that, besides the effect of cell cycle stage, genome-wide spatiotemporal distributions of H3K36me2 and H3K36me3 marks are an additive effort to influence the selection of DSB repair pathway. Our hypothesis is supported by evidence showing that HR is suppressed at the repetitive rDNA locus in yeast, and H3K36me2 promotes NHEJ at unprotected telomeres in mammals (de Krijger *et al*., 2020, Torres-Rosell *et al*., 2007). Meanwhile, Set2 and elongating RNA polymerase II are enriched at DSBs, functioning in DSB repair in the context of transcribed chromatin (Aymard *et al*., 2014, Jha & Strahl, 2014, Tsukuda *et al*., 2005). Additionally, DSBs within actively transcribed genes in which marked with H3K36me3 are targeted to HR repair in a Rad51-dependent manner (Aymard *et al*., 2014). Since the distribution of 20 different histone modifications near DSBs have been characterized across the human genome (Clouaire *et al*., 2018), dissecting individual chromatin signatures would undoubtedly enrich our understanding of how diverse epigenetic modification states on single residue of histones modulate distinct DSB repair pathways.

### Assembly of RPA by H3K36me3 at the distal loci of DSB benefits for tethering of chromatin-centered DDR factors

The RPA complex plays a crucial role in the HR repair network by binding to ssDNA and preserving genome integrity (Pokhrel *et al*., 2019). However, mounting evidence support that DSB repair primarily occurs within the context of chromatin in eukaryotes (Clouaire & Legube, 2015, Smeenk & van Attikum, 2013, Tsukuda *et al*., 2005). Recently, Rtt105 and yBdf1/hTAF1 have been identified as RPA-interacting partners that facilitate RPA loading onto chromatin-based structure (Li *et al*., 2018, Peng *et al*., 2021b). Our study demonstrates that RPA can be recruited to the genome loci through H3K36me3-decorated chromatin, where it may coordinate with various DDR factors to assist in DSB repair. Supporting our result that RPA physically interacts with H3K36me3 nucleosomes, but not H3K36me2 nucleosomes *in vitro* (Figs 3K, 5I, and J). Liu et al. has also shown that RPA directly binds free H3-H4 with a weak affinity, suggesting that RPA-H3-H4 interaction likely occurs on chromatin beyond ssDNA (Liu *et al*., 2017). Compared with the strong ssDNA-bound RPA for protection purpose, weaker chromatin-bound RPA could allow for easier removal of RPA and quick exchange of DDR factors from damaged genomes, thereby facilitating the DSB repair process. The specific mechanism by which H3K36me3-decorated RPA recruitment regulates the exchange of additional effectors to the distal chromatin requires further exploration.

### The conserved role of H3K36me2 and H3K36me3 in DSB repair

Our results demonstrate the conserved function of human partners in specifically recognizing H3K36me2 and H3K36me3-decorated chromatin. Many PWWP domain-containing proteins forms a conserved aromatic cage for methyl-lysine recognition (Qin & Min, 2014). In this study, we reveal that Rfa1 forms a tertiary structure containing a five-β-strands-constituted barrel and a two-helical bundle, which resembles the PWWP domain structures of several known methyl-binding proteins (Appendix Fig S11). Molecular docking indicates these residues form a cage-like structure being sufficient to bind H3K36me3. Crystallography studies in the future may resolve the molecular basis of Rfa1-H3K36me3 recognition. Interestingly, previous studies have shown that the PWWP domain of LEDGF or mismatch recognition protein hMSH6, anchored to H3K36me3-decorated DNA lesions, either facilitating DSB resection or regulating mismatch repair in mammalian cells (Li *et al*., 2013, Pfister *et al*., 2014). The yeast homologs of LEDGF or hMSH6 are not present, but our study highlights the role of H3K36me3 in HR repair and suggests that H3K36me3-decorated chromatin could function as a platform to dynamically associate with different proteins to facilitate DNA repair.

Overall, our study provides a mechanistic insight into how different H3K36 methylation states distinctly function in different DSB repair pathways. These findings would consolidate the notion that the spatiotemporal distributions of distinct H3K36 methylation marks as functional signals to direct the choice of DSB repair pathway for safeguarding genome fidelity.

## Materials and Methods

### Yeast strains

*Saccharomyces cerevisiae* strains were derived from BY4741, JKM139 or tGI354 genetic backgrounds. A complete list of yeast strains is available in Appendix Table S1.

#### Yeast strain construction, cell culture and transformation

Gene disruption and tagging integration were performed using standard PCR-based strategy as described previously (Rothstein, 1983). Corrected strains were verified by colony PCR and/or immunoblotting. The primer sequences used for PCR products are provided in Appendix Table S2.

All yeast cultures were grown at 30 °C. Liquid YP (1% yeast extract, 2% peptone) or SC (6.7 g/L yeast nitrogen base without amino acids, 2 g/L dropout mix) media were supplemented with 2% glucose or other sources of sugar (raffinose or galactose). SC-or YPD-based plates were supplemented with 20 g/L agar. Selection for dominant markers was performed on YPD-based medium supplemented with 200 μg/mL G418 (Beyotime, ST081), 250 μg/mL clonNAT (KLBIOS, PD20181786) or 300 μg/mL hygromycin B (Roche, 10843555001).

For transformation, yeast cultures the logarithmic growth phase were resuspended in LiTE buffer (100 mM LiAc, 10 mM Tris pH 7.5, 1 mM EDTA) and mixed with 10 μL of 10 mg/mL salmon sperm ssDNA, 33.3% [w/v] PEG-3350 and DNA (200 ng of plasmid or purified PCR fragment). Cells were incubated for 30 min at 30 °C, followed by a heat shock at 42 °C for 15 min. Single colonies were isolated from selective media.

### Yeast protein extraction

5 mL mid-log cultures of yeast were quenched in 250 μL of 2 M NaOH with 8% β-mercaptoethanol and incubated on ice for 5 min. Cell pellets were resuspended gently with 250 μL of Buffer A (40 mM HEPES-KOH, pH 7.5, 350 mM NaCl, 0.1% Tween 20, 10% glycerol) followed by 5 min centrifugation at 8000 g. Proteins were resuspended in appropriate volumes of 2 × SDS sample buffer (100 mM Tris–HCl pH 6.8, 2% β-mercaptoethanol, 4% SDS, 0.02% bromophenol blue, 20% glycerol) based on the weight of cell pellets and boiled for 10 min.

### Immunoblotting

Proteins were separated by SDS-PAGE gels, transferred to nitrocellulose membrane (GVS, 1215471), blocked with 3% dry milk dissolved in TBS-T (150 mM NaCl, 20 mM Tris-HCl, 0.05% Tween, pH 7.4), followed by incubation with primary and HRP-coupled secondary antibodies (Jackson Immuno Research Labs, 111-035-003, 115-035-003) diluted in TBS-T. The following commercial primary antibodies were used: anti-H3 (ABclonal, A2348), anti-H3K36me1 (ABclonal, A2364), anti-H3K36me2 (ABclonal, A2365), anti-H3K36me3 (ABclonal, A2366), anti-H2A S129p (Abcam, ab15083), anti-Set2 (Li *et al*., 2017), anti-G6PDH (Sigma, A9521), anti-DYKDDDDK tag (Proteintech, 80010-1-RR), anti-MYC (Proteintech, 60003-2-Ig), anti-GFP (Proteintech, 50430-2-AP), anti-GST(Proteintech, 10000-0-AP), and anti-HA (HuaBio, 0906-1). Signals were revealed with super ECL detection reagents (Yeasen Biotech, 36208ES76); and images were taken by Super ECL Detection Reagent (Yeasen Biotech Co. Ltd, Shanghai, China), and images were taken by Chemiluminescence imaging system (Clinx Science Instruments, Shanghai, China).

### HO-induced DNA damage

Yeast cells were grown in YPD, SC, or SC dropout medium at 30 °C overnight. Cell cultures were diluted to OD_600_ = 0.2, and continued to grow until logarithmic growth phase (OD_600_ ≈ 0.8), unless otherwise indicated. For galactose induction, mid-log phase yeast cells were cultured into YP or SC medium with addition of 2% galactose for the indicated time points.

### Spotting assays

Log phase yeast cells were diluted to OD_600_ = 0.5, five 5× serial dilutions were prepared in sterile microtubes and spotted on SC-based selective medium with bleomycin, camptothecin (CPT), and Etoposide (VP-16), or in the presence of either glucose, raffinose, or galactose as carbon source with the indicated concentrations. Plates were incubated at 30 °C and imaged at 2 – 5 days.

### DSB-induced cell survival assay and quantification

Log-phase yeast cells were grown in the pre-induction medium (SC-Leu + 2% Raffinose) were continued to grow in SC-Leu + 2% Gal medium to induce DSB. At the indicated time points, equal amounts of cells were spotted onto SC-Leu plates. After incubation for several days, colony numbers were counted. Survival rate was calculated using the following formula: (number of colonies grown on SC-Leu plates at the indicated time points / (number of colonies grown on SC-Leu plates at “0 h”) × 100%. At least three independent experiments were performed for each strain.

### Analysis of cleavage efficiency by qRT-PCR

To measure the NHEJ efficiency, HO cleavage analysis was performed as previously described (Gnugge *et al*., 2018). 25 mL-overnight cultured yeast cells in the pre-induction medium YP + Raffinose were cultured to log phase, and then switched to YP medium containing 2% galactose to induce HO cleavage for 1.5 h. After washing with sterile water, cells were resuspended in YPD medium. At each indicated time point, 1 mL recovery cells were collected and genomic DNA was extracted via phenol/chloroform method. Dissolved DNA was used to measure HO cleavage levels by measuring the amounts of induced samples using PCR. The values in samples with time starting damage recovery was set up as “1”, and relative values of induced samples at each time point were normalized to that at time 0. Three independent experiments were performed for each strain.

### Fluorescence microscopy

HO-induced DSB was activated by adding of 2% galactose to the log phase (1×10^7^/mL) yeast cells grown in the pre-induction medium (YP-Raffinose). Samples were collected at the indicated time points after DSB induction and then were spread on microslides. Bright field and fluorescent images were captured under a confocal laser scanning microscope (TCS SP8 MP, Leica, Germany). The percentage of cells carrying Rad51-GFP foci was calculated after analyzing three independent experiments. Approximately 100 cells were counted from each experiment.

### Analysis of DSB end resection

To Monitor 5’-end resection assays, quantitative PCR (qPCR) or Southern blot methods were employed with some modifications (Gnugge *et al*., 2018, Peng *et al*., 2021a). DSB was induced by adding 2% galactose to logarithmic phase cells grown in the pre-induction medium (YP + Raffinose). Samples were collected at the indicated time points after DSB induction. Genomic DNA was extracted using phenol/chloroform and dissolved in 100 μL sterile water.

For qPCR approach, 10 μg genomic DNA was digested with 20 units *Sty* I (NEB, R3500S) at 37 °C for 3 h. qPCR was performed using MonAmpTM SYBR Green quantitative qPCR Mix (Monad, MQ10101S). A three-step PCR program was used according to the manufacturer’s protocol. The total amounts of DNA at three sites distal to the break were determined by qPCR and normalized to an amplicon in the *ACT* gene to account for differences in template concentrations. As the 5’ strand undergoes degradation at DSB ends, the restriction enzyme is unable to cleave ssDNA, allowing the fraction of DNA to be PCR amplified. The qPCR primer sequences were listed in Appendix Table S3.

To measure resection kinetics examined by Southern blot, 20 μg purified genomic DNA was digested with 30 units *EcoR* I (NEB, R3101L) and 30 units *Hind* III (NEB, R3104V) followed by resolved on a 0.8% agarose gel. The restricted DNA fragments were transferred onto a positively charged Nylon membrane (GE Healthcare, RPN303B) and Southern blotting was performed to measure the percentage of unresected DNA. Probe preparation and subsequent DIG detection were conducted using the DIG Northern Starter Kit (Roche, 12039672910) according to the manufacturer’s instruction. The band density corresponding to each probe over time is quantified by ImageJ to visualize the resection kinetics (“*TRA1*” signals served as normalized controls).

### Plasmid-based DSB re-ligation assay

The plasmid recombination assay was performed as previously described (Jessulat *et al*., 2008). Briefly, 0.9 μg of linearized pRS416 digested with 10 units *EcoR* I and 100 ng of circular pRS415 plasmids (as a normalized control) were co-transformed into the indicated strains. Transformations were spotted onto SC-Ura and SC-Leu plates. The colonies were counted after 4 days. NHEJ efficiency was calculated by the ratio of uracil prototrophic growing colonies relative to leucine prototrophic growing colonies. Three independent transformation reactions were performed for each strain.

### Integration rate assay

The integration rate assay was performed as described previously (Chen *et al*., 2012). To measure the HR efficiency of different strains, 300 ng of *ADH4Δ*::*URA3* gene knockout cassettes, containing ∼1.5 kb of flanking sequence on each side of the endogenous *ADH4* locus or 50 ng of pRS416 (as a normalized control) were transformed into the indicated strains, respectively. Individual transformation was plated onto SC-Ura plates. Integration rate was calculated by the ratio of growing colonies transformed with gene knockout cassettes relative to growing colonies transformed with pRS416. Three independent transformation reactions were performed for each strain.

### Protein expression and purification

Full-length and truncated proteins were amplified from cDNA derived from human or yeast cells, and PCR products were subcloned into pGEX-6p-1 vector using one-step cloning strategy according to the protocols of the ClonExpress II One Step Cloning Kit (Vazyme, C112-01). Site-directed point mutations were generated using the site-directed mutagenesis protocol (Stratagene), and sequences were confirmed by DNA sequencing. Recombination proteins were induced 0.2 mM isopropyl β-D-1-thiogalactopyranoside (IPTG) in Escherichia coli BL21 (DE3) at OD600 = 0.8 and chaperone protein TIG was induced by the addition of 0.5 mg/mL L-arabinose (Sangon, A610071-0025) stem from plasmid pTf16, which can increase the solubility of target protein. Cells were cultured at 20 °C for 6 h before harvest. Cells were lysed in GST lysis buffer (50 mM Tris-HCl, pH 7.5, 150 mM NaCl, 0.05% NP-40) by sonication. The lysate was clarified by centrifugation at 12,000 rpm for 30 min at 4 °C. Appropriate glutathione Sepharose beads (Smart lifesciences, SA010025) were pre-washed with PBS and equilibrated in GST lysis buffer. Beads were incubated with cell lysates at 4 °C for 4 h with rotation, and beads containing bound proteins were washed 3 times with PBS and proteins were eluted with elution buffer (50 mM Tris-HCl, pH 7.5, 20 mM reduced glutathione). Protein concentration were estimated by SDS-PAGE using BSA as a standard.

### GST pull-down assay

The recombinant GST proteins immobilized on glutathione agarose beads was incubated with 100 μg histone peptides at 4 °C for 4 h with continuous rotation. Beads were washed extensively 3 times with wash buffer (20 mM Tris-HCl, pH 7.4, 200 mM NaCl, 0. 5 mM EDTA, and 10% glycerol). Bound peptides were eluted by boiling beads in 2× SDS loading buffer (without bromophenol blue) for 4 min after extensively washing with lysis buffer. 25% input samples were spotted on NC membrane and bound peptides were detected by dot blotting.

### Peptide pull-down assay

100 μg of biotinylated histone peptides with different H3K36 modifications were incubated with 20 μg of GST-fused proteins in binding buffer (50 mM Tris-HCl pH 7.5, 200 mM NaCl, 0.05% NP-40, 1 mM PMSF) overnight. Streptavidin beads (Smart lifesciences, SA092005)) were added to the mixture, and then continuously incubated for 1 h with rotation. After extensively washing with lysis buffer for 3 times, bound proteins were eluted by boiling beads in 2× SDS loading buffer for 4 min. Elutes were loaded onto SDS-PAGE and signals were examined by Western blotting. The peptide sequences were listed in the Appendix Table S5.

### Co-immunoprecipitation

50 mL of yeast cells culture (OD_600_ ≈ 0.8) were collected and resupended in 500 μL of lysis buffer (25 mM Tris–HCl pH 8.8, 300 mM NaCl, 5% glycerol, 0.2% Triton X-100, 1 mM PMSF), mixed with 300 μL of glass beads and homogenized by bead beater, 8× 30 s cycles of full speed, cooling samples on ice for at least 1 min in every 4 cycles. All subsequent procedures were performed at 4 °C. The extracts were clarified by centrifugation at 12000 g for 10 min, 20 μL input was collected and the remaining lysates were incubated with 20 μL 50% slurry of anti-Flag agarose beads (Smart lifesciences, SA042C), anti-HA agarose beads (AOGMA, AGM90054) or protein A/G beads with α-MYC antibody (Proteintech, 60003-2-Ig) for 6 h. Beads were washed 3 times with lysis buffer and boiled in 20 μL of 2× SDS sample buffer. Input and IP samples were subjected to immunoblotting.

### Chromatin immunoprecipitation (ChIP)

ChIP was carried out as previously described with minor modifications (Shu et al, 2020a). Briefly, cells were grown to OD600 ≈ 0.8, and were cross-linked with 1% formaldehyde for 13 min and quenched with 0.1 M glycine for 5 min. Frozen pellets were lysed in 0.25 mL ChIP lysis buffer A. After glass-bead beating, chromatin was sonicated for 24 cycles (high output, 30 s-on, 30 s-off) using the Bioruptor Plus (Diagenode, Belgium) to produce fragments of ∼ 250 bp. Protein concentration was measured and equivalent quantities of total protein (Input) were used for chromatin immunoprecipitation. Lysates were incubated with 10 μL-□Flag or IgG beads at 4 °C overnight with continuous rotation. For H3/H3K36me2/H3K36me3 ChIP, Iysates were incubated with 2 □g appropriate antibodies at 4 °C overnight with rotation, 10 μL volume of protein G beads were then added for extra 1 h. Beads were sequentially washed once with ChIP lysis buffer A for 10 min, once with ChIP lysis buffer B for 10 min, and once with LiCl/NP40 buffer for 10 min. Bound chromatin fractions were eluted from beads by two sequential incubations with 200 μL of elution buffer at room temperature for 15 min each. After de-crosslinking by incubation elutes at 65°C for > 5 h, 20□g proteinase K (ABclonal, RP02503) were added into each sample and incubated at 42°C for 1 h. DNA was purified by universal DNA purification kit (Tiangen, DP314-03) according to manufacturer’s protocol. DNA was eluted from the column with 30□□L sterile H2O. qPCRs were performed using MonAmpTM SYBR green quantitative qPCR mix with oligonucleotide pairs listed in the Appendix Table S3.

### Microscale Thermophoresis (MST)

MST experiments were performed on a Monolith NT.115 instrument (NanoTemper Technologies, Germany), using standard treated capillaries (Nanotemper). Purified GST fusion protein were fluorescently labeled using the Monolith series Protein Labeling Kit RED-NHS 2nd Generation (Nanotemper, MO-L011) according to manufacturer’s instructions. For the MST assay, 100 nM fluorescently labeled GST-Rfa1 or GST-Ku70 were mixed with series of two-fold diluted native nucleosomes in MST binding buffer (20 mM Tris-HCl pH 8.0, 50 mM NaCl, 0.05% Tween-20). The measurements were performed at 26 °C using 20% LED power and 40% MST power, with 20 s laser-on time and 5 s laser-off time. Thermophoresis data were analyzed by MO. Affinity Analysis software (NanoTemper) using the data on time 1.5 s. At least two independent experiments were performed for each measurement.

### Mammalian cell culture, transfection, plasmid construction and protein extraction

HEK293T cells (ATCC, CRL-11268) and U2OS cells (ATCC, HTB-96) were cultured in DMEM medium (Monad, CC00101S) supplemented with 10% fetal bovine serum (FBS, Lonsera, S711-001S) and penicillin/streptomycin (BasalMedia, S110JV) at 37 °C. When the cell density reached to 70% confluency, plasmids were transfected into cell culture using lipofectamine 2000 (Invitrogen, 11668019) following the manufacturer’s instruction. Cells were collected after 48 h-72 h.

Human full-length RPA1/KU70 was amplified from cDNA derived from human A549 cells and then was subcloned into a pCS2-3× Flag vector using one-step cloning strategy according to the protocols of the ClonExpress II One Step Cloning Kit (Vazyme, C112-01). Site-directed point mutations were generated using the Site-Directed Mutagenesis protocol (Stratagene), and sequences were confirmed by DNA sequencing. The PCR primer sequences used to generate these constructs or mutations were listed in Appendix Table S2, and plasmids used in this study were listed in Appendix Table S4.

For protein extraction, cells were firstly dissolved in lysis buffer (25 mM Tris–HCl pH 8.8, 300 mM NaCl, 5% glycerol, 0.2% Triton X-100, 1 mM PMSF) and then placed on ice for 30 min, followed by mild sonication.

### Quantification and statistical analysis

Unless otherwise stated, Western blot data were quantified using ImageJ to measure the relative intensity of each band. Quantification data were presented as the mean ± SD (standard deviation) from at least three independent experiments using GraphPad Prism 9.5.2. Statistical differences were determined by two-tailed unpaired *t*-test, and a *P*-value of less than 0.05, 0.01, 0.001 or 0.0001 was considered statistically significant and marked as ‘*’, ‘**’, ‘***’, ‘****’, respectively. ‘ns’ indicates ‘not significant’.

## Data availability

The datasets produced in this study are available in the following database: Imaging and digital dataset: Image Data Resource DOI:10.17632/nr9sx8by4f.1

## Acknowledgments

We thank Dr. Grzegorz Ira (Baylor College of Medicine) for tGI354 strain, Dr. Qiang Chen (Wuhan University) for U2OS cell line, Dr. Zheng Zhou (Institute of Biophysics, CAS), Mr. Jiangpeng Feng, and Mr. Yang Li (Wuhan University) for technique assistance. This work was supported by the National Key R&D program of China (2019YFA0802501), the National Natural Science Foundation of China (32270617, 31971231), the Fundamental Research Funds for the Central Universities (2042022dx0003), and the Application Fundamental Frontier Foundation of Wuhan (2020020601012225).

## Author Contributions

H.-N.D. conceived the project, R.C., M.-J.Z., Y.-M.L., and R.-X.W. conducted most experiments, R.C., M.-J.Z., Y.-M.L., A.-H.L., Y.-C.M, and H.-N.D. acquired and analyzed data. X.C. provided technical support and research resources. R.C., M.-J.Z., and H.-N.D. with the help of X.C. wrote the manuscript.

## Disclosure statement and competing interests

The authors declare that they have no conflict of interest.

## References

Aymard F, Bugler B, Schmidt CK, Guillou E, Caron P, Briois S, Iacovoni JS, Daburon V, Miller KM, Jackson SP et al. (2014) Transcriptionally active chromatin recruits homologous recombination at DNA double-strand breaks. Nat Struct Mol Biol 21: 366–74

Bird AW, Yu DY, Pray-Grant MG, Qiu Q, Harmon KE, Megee PC, Grant PA, Smith MM, Christman MF (2002) Acetylation of histone H4 by Esa1 is required for DNA double-strand break repair. Nature 419: 411–5

Bunting SF, Callen E, Wong N, Chen HT, Polato F, Gunn A, Bothmer A, Feldhahn N, Fernandez-Capetillo O, Cao L et al. (2010) 53BP1 inhibits homologous recombination in Brca1-deficient cells by blocking resection of DNA breaks. Cell 141: 243–54

Carvalho S, Vitor AC, Sridhara SC, Martins FB, Raposo AC, Desterro JM, Ferreira J, de Almeida SF (2014) SETD2 is required for DNA double-strand break repair and activation of the p53-mediated checkpoint. Elife 3: e02482

Chen X, Cui D, Papusha A, Zhang X, Chu CD, Tang J, Chen K, Pan X, Ira G (2012) The Fun30 nucleosome remodeller promotes resection of DNA double-strand break ends. Nature 489: 576–80

Chiruvella KK, Liang Z, Wilson TE (2013) Repair of double-strand breaks by end joining. Cold Spring Harb Perspect Biol 5: a012757

Clouaire T, Legube G (2015) DNA double strand break repair pathway choice: a chromatin based decision? Nucleus 6: 107–13

Clouaire T, Rocher V, Lashgari A, Arnould C, Aguirrebengoa M, Biernacka A, Skrzypczak M, Aymard F, Fongang B, Dojer N et al. (2018) Comprehensive Mapping of Histone Modifications at DNA Double-Strand Breaks Deciphers Repair Pathway Chromatin Signatures. Mol Cell 72: 250–262 e6

Daley JM, Palmbos PL, Wu D, Wilson TE (2005) Nonhomologous end joining in yeast. Annu Rev Genet 39: 431–51

de Krijger I, van der Torre J, Peuscher MH, Eder M, Jacobs JJL (2020) H3K36 dimethylation by MMSET promotes classical non-homologous end-joining at unprotected telomeres. Oncogene 39: 4814–4827

DiFiore JV, Ptacek TS, Wang Y, Li B, Simon JM, Strahl BD (2020) Unique and Shared Roles for Histone H3K36 Methylation States in Transcription Regulation Functions. Cell Rep 31: 107751

Dronamraju R, Jha DK, Eser U, Adams AT, Dominguez D, Choudhury R, Chiang YC, Rathmell WK, Emanuele MJ, Churchman LS et al. (2018) Set2 methyltransferase facilitates cell cycle progression by maintaining transcriptional fidelity. Nucleic Acids Res 46: 1331–1344

Escribano-Diaz C, Orthwein A, Fradet-Turcotte A, Xing M, Young JT, Tkac J, Cook MA, Rosebrock AP, Munro M, Canny MD et al. (2013) A cell cycle-dependent regulatory circuit composed of 53BP1-RIF1 and BRCA1-CtIP controls DNA repair pathway choice. Mol Cell 49: 872–83

Fell VL, Schild-Poulter C (2015) The Ku heterodimer: function in DNA repair and beyond. Mutat Res Rev Mutat Res 763: 15–29

Findlay S, Heath J, Luo VM, Malina A, Morin T, Coulombe Y, Djerir B, Li Z, Samiei A, Simo-Cheyou E et al. (2018) SHLD2/FAM35A co-operates with REV7 to coordinate DNA double-strand break repair pathway choice. EMBO J 37

Fnu S, Williamson EA, De Haro LP, Brenneman M, Wray J, Shaheen M, Radhakrishnan K, Lee SH, Nickoloff JA, Hromas R (2011) Methylation of histone H3 lysine 36 enhances DNA repair by nonhomologous end-joining. Proc Natl Acad Sci U S A 108: 540–5

Gilbert TM, McDaniel SL, Byrum SD, Cades JA, Dancy BC, Wade H, Tackett AJ, Strahl BD, Taverna SD (2014) A PWWP domain-containing protein targets the NuA3 acetyltransferase complex via histone H3 lysine 36 trimethylation to coordinate transcriptional elongation at coding regions. Mol Cell Proteomics 13: 2883–95

Gnugge R, Oh J, Symington LS (2018) Processing of DNA Double-Strand Breaks in Yeast. Methods Enzymol 600: 1–24

Gopalakrishnan R, Marr SK, Kingston RE, Winston F (2019) A conserved genetic interaction between Spt6 and Set2 regulates H3K36 methylation. Nucleic Acids Res 47: 3888–3903

Hu M, Sun XJ, Zhang YL, Kuang Y, Hu CQ, Wu WL, Shen SH, Du TT, Li H, He F et al. (2010) Histone H3 lysine 36 methyltransferase Hypb/Setd2 is required for embryonic vascular remodeling. Proc Natl Acad Sci U S A 107: 2956–61

Husmann D, Gozani O (2019) Histone lysine methyltransferases in biology and disease. Nat Struct Mol Biol 26: 880–889

Iacovoni JS, Caron P, Lassadi I, Nicolas E, Massip L, Trouche D, Legube G (2010) High-resolution profiling of gammaH2AX around DNA double strand breaks in the mammalian genome. EMBO J 29: 1446–57

Ira G, Haber JE (2002) Characterization of RAD51-independent break-induced replication that acts preferentially with short homologous sequences. Mol Cell Biol 22: 6384–92

Jasin M, Rothstein R (2013) Repair of strand breaks by homologous recombination. Cold Spring Harb Perspect Biol 5: a012740

Jessulat M, Alamgir M, Salsali H, Greenblatt J, Xu J, Golshani A (2008) Interacting proteins Rtt109 and Vps75 affect the efficiency of non-homologous end-joining in Saccharomyces cerevisiae. Arch Biochem Biophys 469: 157–64

Jha DK, Strahl BD (2014) An RNA polymerase II-coupled function for histone H3K36 methylation in checkpoint activation and DSB repair. Nat Commun 5:3965

Kaur E, Nair J, Ghorai A, Mishra SV, Achareker A, Ketkar M, Sarkar D, Salunkhe S, Rajendra J, Gardi N et al. (2020) Inhibition of SETMAR-H3K36me2-NHEJ repair axis in residual disease cells prevents glioblastoma recurrence. Neuro Oncol 22: 1785–1796

Kockler ZW, Osia B, Lee R, Musmaker K, Malkova A (2021) Repair of DNA Breaks by Break-Induced Replication. Annu Rev Biochem 90: 165–191

Kowalczykowski SC (2015) An Overview of the Molecular Mechanisms of Recombinational DNA Repair. Cold Spring Harb Perspect Biol 7

Li F, Mao G, Tong D, Huang J, Gu L, Yang W, Li GM (2013) The histone mark H3K36me3 regulates human DNA mismatch repair through its interaction with MutSalpha. Cell 153: 590–600

Li F, Zheng LD, Chen X, Zhao X, Briggs SD, Du HN (2017) Gcn5-mediated Rph1 acetylation regulates its autophagic degradation under DNA damage stress. Nucleic Acids Res 45: 5183–5197

Li J, Sun H, Huang Y, Wang Y, Liu Y, Chen X (2019) Pathways and assays for DNA double-strand break repair by homologous recombination. Acta Biochim Biophys Sin (Shanghai) 51: 879–889

Li S, Xu Z, Xu J, Zuo L, Yu C, Zheng P, Gan H, Wang X, Li L, Sharma S et al. (2018) Rtt105 functions as a chaperone for replication protein A to preserve genome stability. EMBO J 37

Liu S, Xu Z, Leng H, Zheng P, Yang J, Chen K, Feng J, Li Q (2017) RPA binds histone H3-H4 and functions in DNA replication-coupled nucleosome assembly. Science 355: 415–420

Lu Z, Lin Z (2019) Pervasive and dynamic transcription initiation in Saccharomyces cerevisiae. Genome Res 29: 1198–1210

Marechal A, Zou L (2015) RPA-coated single-stranded DNA as a platform for post-translational modifications in the DNA damage response. Cell Res 25: 9–23

McDaniel SL, Strahl BD (2017) Shaping the cellular landscape with Set2/SETD2 methylation. Cell Mol Life Sci 74: 3317–3334

Mei YC, Feng J, He F, Li YM, Liu Y, Li F, Chen Y, Du HN (2021) Set2-mediated H3K36 methylation states redundantly repress the production of antisense transcripts: role in transcription regulation. FEBS Open Bio 11: 2225–2235.

Miyazaki T, Bressan DA, Shinohara M, Haber JE, Shinohara A (2004) In vivo assembly and disassembly of Rad51 and Rad52 complexes during double-strand break repair. EMBO J 23: 939–49

Pai CC, Deegan RS, Subramanian L, Gal C, Sarkar S, Blaikley EJ, Walker C, Hulme L, Bernhard E, Codlin S et al. (2014) A histone H3K36 chromatin switch coordinates DNA double-strand break repair pathway choice. Nat Commun 5: 4091

Pai CC, Kishkevich A, Deegan RS, Keszthelyi A, Folkes L, Kearsey SE, De Leon N, Soriano I, de Bruin RAM, Carr AM et al. (2017) Set2 Methyltransferase Facilitates DNA Replication and Promotes Genotoxic Stress Responses through MBF-Dependent Transcription. Cell Rep 20: 2693–2705

Peng H, Zhang S, Chen X (2021a) Monitoring 5’-End Resection at Site-Specific Double-Strand Breaks by Southern Blot Analysis. Methods Mol Biol 2196: 245–255

Peng H, Zhang S, Peng Y, Zhu S, Zhao X, Zhao X, Yang S, Liu G, Dong Y, Gan X et al. (2021b) Yeast Bromodomain Factor 1 and Its Human Homolog TAF1 Play Conserved Roles in Promoting Homologous Recombination. Adv Sci (Weinh) 8: e2100753

Pfister SX, Ahrabi S, Zalmas LP, Sarkar S, Aymard F, Bachrati CZ, Helleday T, Legube G, La Thangue NB, Porter AC et al. (2014) SETD2-dependent histone H3K36 trimethylation is required for homologous recombination repair and genome stability. Cell Rep 7: 2006–18

Pokhrel N, Caldwell CC, Corless EI, Tillison EA, Tibbs J, Jocic N, Tabei SMA, Wold MS, Spies M, Antony E (2019) Dynamics and selective remodeling of the DNA-binding domains of RPA. Nat Struct Mol Biol 26: 129–136

Qin S, Min J (2014) Structure and function of the nucleosome-binding PWWP domain. Trends Biochem Sci 39: 536–47

Rothstein RJ (1983) One-step gene disruption in yeast. Methods Enzymol 101: 202–11

Scully R, Panday A, Elango R, Willis NA (2019) DNA double-strand break repair-pathway choice in somatic mammalian cells. Nat Rev Mol Cell Biol 20: 698–714

Sengupta D, Zeng L, Li Y, Hausmann S, Ghosh D, Yuan G, Nguyen TN, Lyu R, Caporicci M, Morales Benitez A et al. (2021) NSD2 dimethylation at H3K36 promotes lung adenocarcinoma pathogenesis. Mol Cell 81: 4481–4492 e9

Shirane K, Miura F, Ito T, Lorincz MC (2020) NSD1-deposited H3K36me2 directs de novo methylation in the mouse male germline and counteracts Polycomb-associated silencing. Nat Genet 52: 1088–1098

Smeenk G, van Attikum H (2013) The chromatin response to DNA breaks: leaving a mark on genome integrity. Annu Rev Biochem 82: 55–80

Symington LS (2014) End resection at double-strand breaks: mechanism and regulation. Cold Spring Harb Perspect Biol 6

Symington LS, Gautier J (2011) Double-strand break end resection and repair pathway choice. Annu Rev Genet 45: 247–71

Torres-Rosell J, Sunjevaric I, De Piccoli G, Sacher M, Eckert-Boulet N, Reid R, Jentsch S, Rothstein R, Aragon L, Lisby M (2007) The Smc5-Smc6 complex and SUMO modification of Rad52 regulates recombinational repair at the ribosomal gene locus. Nat Cell Biol 9: 923–31

Tsukuda T, Fleming AB, Nickoloff JA, Osley MA (2005) Chromatin remodelling at a DNA double-strand break site in Saccharomyces cerevisiae. Nature 438: 379–83

Wagner EJ, Carpenter PB (2012) Understanding the language of Lys36 methylation at histone H3. Nat Rev Mol Cell Biol 13: 115–26

Waterman DP, Zhou F, Li K, Lee CS, Tsabar M, Eapen VV, Mazzella A, Haber JE (2019) Live cell monitoring of double strand breaks in S. cerevisiae. PLoS Genet 15: e1008001

Weinberg DN, Papillon-Cavanagh S, Chen H, Yue Y, Chen X, Rajagopalan KN, Horth C, McGuire JT, Xu X, Nikbakht H et al. (2019) The histone mark H3K36me2 recruits DNMT3A and shapes the intergenic DNA methylation landscape. Nature 573: 281–286

Yano S, Ishiuchi T, Abe S, Namekawa SH, Huang G, Ogawa Y, Sasaki H (2022) Histone H3K36me2 and H3K36me3 form a chromatin platform essential for DNMT3A-dependent DNA methylation in mouse oocytes. Nat Commun 13: 4440

Zhang Y, Hefferin ML, Chen L, Shim EY, Tseng HM, Kwon Y, Sung P, Lee SE, Tomkinson AE (2007) Role of Dnl4-Lif1 in nonhomologous end-joining repair complex assembly and suppression of homologous recombination. Nat Struct Mol Biol 14: 639–46

Zheng S, Li D, Lu Z, Liu G, Wang M, Xing P, Wang M, Dong Y, Wang X, Li J et al. (2018) Bre1-dependent H2B ubiquitination promotes homologous recombination by stimulating histone eviction at DNA breaks. Nucleic Acids Res 46: 11326–11339

Zhu Z, Chung WH, Shim EY, Lee SE, Ira G (2008) Sgs1 helicase and two nucleases Dna2 and Exo1 resect DNA double-strand break ends. Cell 134: 981–94

